# Hepatic aldose reductase drives a Warburg effect-like metabolic reprogramming to promote insulin resistance, fatty liver and obesity

**DOI:** 10.1101/2024.09.23.614395

**Authors:** Dan Song, Dianqiang Yang, Longping Wen, Feng Zheng, James Y. Yang

## Abstract

**Background & Aims:** Emerging evidence suggest that abnormal activation of aldose reductase/the polyol pathway (Ar/PP) is associated with the pathogenesis or development of fatty liver, obesity and metabolic syndrome. However, the underlying mechanisms were unclear. In this study, we investigated the metabolic reprogramming following activation or inhibition of Ar, the first and the rate-limiting enzyme of PP. We also investigated the long-term effects of Ar/PP-mediated metabolic shift *in vivo*.

**Methods:** Metabolomic analyses were performed with the AB-SCIE QTRAP-5500 LC-MS/MS System for control mouse hepatocytes and hepatocytes stably overexpressing Ar and exposed to 25 mM glucose. Glycolysis stress tests and mitochondrial stress tests were performed using the Seahorse Bioscience Extracellular Flux Analyzer. The *in vivo* long-term effects of Ar overexpression and inhibition were evaluated in either transgenic mice overexpressing AR or a line of double transgenic mice carrying an Ar-null mutation and an Agouti-yellow *A^y^* mutation.

**Results:** Abnormal activation of Ar in hepatocytes was found to trigger and drive a drastic Warburg effect-like metabolic reprogramming, induce *de novo* lipogenesis, and alter insulin and AMP-activated protein kinase signaling. In glucose-fed *AR*-overexpressing transgenic mice, AR activation causes systemic alterations in physiological parameters and the development of overt phenotypes of insulin resistance, fatty liver, obesity. In the yellow obese syndrome mice, *Ar* deficiency greatly improves *Agouti A^y^* mutation-induced abnormalities.

**Conclusions:** Collectively, the results highlight the important contribution of Ar/PP or the putative pseudo-glycolysis in hepatic metabolic homeostasis and the development of metabolic diseases. These findings have profound implications for the development of therapeutic strategies or drugs against metabolic diseases and cancer.

**GRAPHICAL ABSTRACT:** 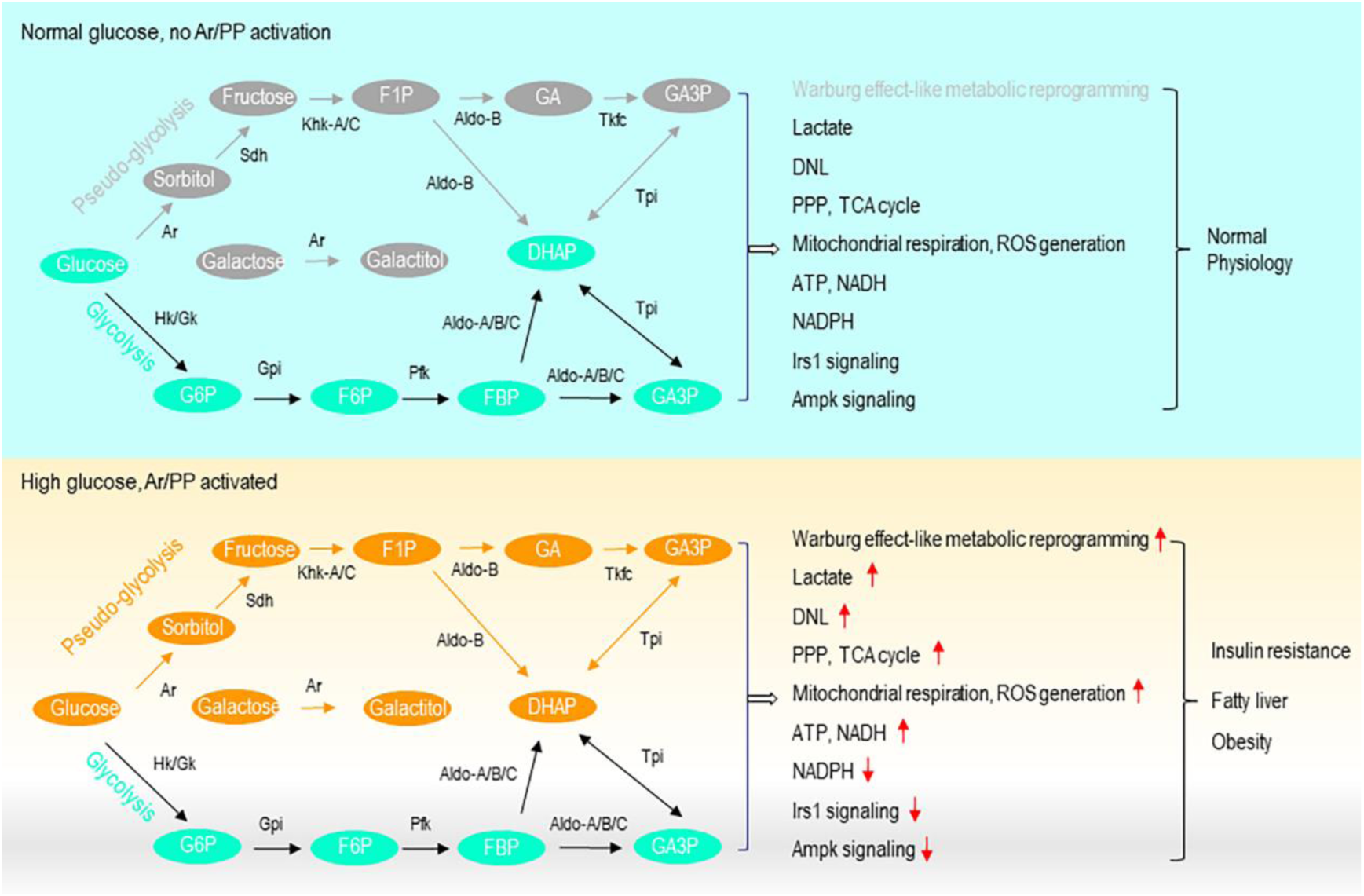

**Highlights:**

- Activation of aldose reductase triggers and drives a Warburg effect-like metabolic eprogramming in hepatocytes.
- Liver-specific activation of the polyol pathway leads to insulin resistance, fatty liver and obesity.
- Inhibition of aldose reductase greatly ameliorates *Agouti A^y^*-induced metabolic abnormalities.

**Impact and implications:** This study reveals that abnormal activation of Ar/PP will trigger and drive a Warburg effect-like metabolic reprogramming in hepatocytes. In normal subjects, Ar/PP mediated metabolic reprogramming tends to promote lipogenesis, insulin resistance, fatty liver and obesity. In cancer cells, Ar/PP mediated metabolic reprogramming will be part of the Warburg effect to support the growth and proliferation of cancer cells. These findings imply that Ar and its down-stream metabolic enzymes are important therapeutic targets for cancers and metabolic diseases.

## INTRODUCTION

Type II diabetes, non-alcoholic fatty liver disease (NAFLD) and non-alcoholic steatohepatitis (NASH), obesity and metabolic syndrome are the major health challenges of our modern societies. During the last few decades, a rapid increase in carbohydrate consumption in the developed countries is found to coincide with a dramatic rise of obesity and diabetes and cardiovascular diseases, raising concerns about the negative health impact of the overconsumption of added sugars (1–5). Fructose is an isomer of glucose. However, fructose is almost twice as sweet as glucose and therefore is widely used (especially in the form of high fructose corn syrup) as a sweetener in processed food, deserts, candies, drinks or beverages. In hepatocytes, a recent study shows that the rate of fructose oxidation at near physiological concentration (1 mM) exceeds that of 25 mM glucose (6), suggesting distinct metabolic efficiency and fate. Meanwhile, a large collection of clinical and experimental evidence overwhelmingly suggests that added fructose tends to induce insulin resistance, glucose intolerance, hyperinsulinemia, hyperleptinemia, endoplasmic reticulum stress, inflammation, dyslipidemia and *de novo* lipogenesis (DNL) but tends to suppress β-oxidation of fatty acid (6–9). Fructose is thus regarded as the principal driving force behind diabetes, NAFLD, NASH, obesity, metabolic syndrome, cardiovascular diseases and cancer (10–18).

Dietary ingestion, however, is not the only source for fructose. In fact, fructose can be produced from glucose *in vivo* by a glucose-metabolic shunt called the polyol pathway (PP) (10,19). In the first step of PP, aldose reductase (Ar) catalyzes the conversion of glucose to sorbitol, with the aid of its co-factor NADPH (Fig. S1). In the second step of PP, sorbitol is oxidized by sorbitol dehydrogenase (Sdh), using NAD^+^ as a cofactor, to form fructose and NADH. The reaction catalyzed by Ar is the rate-limiting step of PP. Ar protein, however, has a low affinity for glucose, with a *Km* between 30-80 mM (20,21). This implies that PP operates only when glucose is abundant. Although it has been shown recently that the intestine and kidney might also contribute (22–24), the liver is the most import tissue for fructose metabolism. Regardless of exogenous or endogenous sources, in the liver fructose is metabolized though fructolysis (6,7,10,17). In the first step of fructolysis, fructose is phosphorylated by ketohexokinase (Khk) to form fructose 1-phosphate (F1P). In the second step of fructolysis, the six-carbon F1P is split by aldolase B (AldoB) into two three-carbon trioses, namely glyceraldehyde and dihydroxyacetone phosphate (DHAP). In an extra step of fructolysis, glyceraldehyde is further converted to glyceraldehyde 3-phosphate (GA3P) by triose kinase/FMN cyclase (Tkfc) (25). From DHAP and GA3P on up to the formation of pyruvate, the remaining 6 steps of fructolysis are exactly the same as that of the conventional glycolysis, using exactly the same liver metabolic enzymes. To consider the endogenous fructose production and utilization together, we use hereafter a term “pseudo-glycolysis” to refer to the whole process through which glucose is converted into fructose by PP and fructose is metabolized through fructolysis down to the formation of two molecules of pyruvate.

The putative PP embedded pseudo-glycolysis and the conventional glycolysis both starts with glucose and ends up with pyruvate (Fig. S1). When a molecule of glucose is catabolized through the conventional glycolysis to form 2 molecules of pyruvate, it uses 2 molecules of ATP in the priming reactions and yields 4 molecules of ATP and 2 molecules of NADH in the pay-off phase. The pseudo-glycolysis also uses 2 molecules of ATP in the priming reactions and yields 4 molecules of ATP later. However, the putative pseudo-glycolysis generates 3 molecules of NADH, as compared to 2 molecules in the conventional glycolysis, although at a cost of the consumption of 1 molecule of NADPH. In terms of ATP production, the putative pseudo-glycolysis will probably be more efficient than the conventional glycolysis as 1 molecule of NADH in the mitochondria is equivalent to 2-3 molecules of ATP. A single most important difference between the conventional glycolysis and the putative pseudo-glycolysis, nevertheless, is that the formation of F1P from fructose in the pseudo-glycolysis completely bypasses the major regulatory step as seen in the conventional glycolysis, *i.e.*, the phosphorylation of fructose 6-phosphoate to form fructose-1,6-bisphosphate (FBP) catalyzed by phosphofructokinase (Pfk) (26–28). Pfk is strongly feedback-inhibited by ATP, citrate, low pH and oxygen (27), so the flux of the conventional glycolysis is often self-limited. Since the putative pseudo-glycolysis lacks this feedback inhibition, it can be expected that a large amount of glucose can be fluxed through this pathway to produce intermediates used in other metabolic pathways including glycolysis, pentose phosphate pathway, citric acid cycle (TCA), oxidative phosphorylation, lipogenesis, gluconeogenesis/glycogenesis, according to the needs of the cells. It is well established that Ar/PP plays important roles in the pathophysiology of diabetic complications (21,29). The physiological roles of PP/the putative pseudo-glycolysis, however, remain to be fully investigated. Recently, evidence has emerged showing that fructose produced endogenously by PP might also contribute significantly to the pathogenesis of diseases including diabetes-associated dyslipidemia (30), NAFLD (31), nonalcoholic steatohepatitis (NASH) (32), alcoholic steatosis (AFLD) (33,34), fatty liver, obesity and metabolic syndrome (19,35,36) and hepatocellular carcinoma (HCC) (37,38) and gastric cancer (39). In these studies, the flux of glucose through PP/the putative pseudo-glycolysis was usually blocked using genetically deleted *Ar* or *Khk* mutant mice. Alternatively, Ar inhibitor (ARI) was used to inhibit the first and the rate-limiting step of PP. Results from these studies clearly show that deficiency/inhibition of Ar or the lack of Khk significantly improve dyslipidemia, AFLD, NAFLD, NASH, fatty liver, obesity and HCC. More importantly, however, is how the overactivation of Ar/PP or the putative pseudo-glycolysis might trigger and drive the overall metabolic dysregulation, which might not simply be the opposite to the inhibition of Ar or the blockade of PP. The mechanisms underlying Ar/PP-induced metabolic dysfunction thus remain to be fully elucidated.

With metabolomic analyses, we found in this study that hepatic Ar activation caused a drastic Warburg Effect-like metabolic reprogramming in cultured AML12 hepatocytes, affecting predominantly the metabolism of carbohydrates, lipids, pyrimidines and purines. In terms of metabolic signaling, we found that Ar overexpression significantly suppressed whereas Ar knockdown greatly increased the expression of insulin receptor substrate-1 (Irs1). Meanwhile, Ar overexpression significantly suppressed whereas Ar knockdown greatly increased the phosphorylation of liver kinase B1 (Lkb1), AMP-activated protein kinase-α (Ampkα) and acetyl-CoA carboxylase (Acc). Lkb1 is the upstream activator of Ampk and Acc is one of the downstream targets of Ampk and the rate-limiting enzyme for lipid synthesis. *In vivo* in *Ar*-overexpressing transgenic mice, Ar activation -mediated suppression of Irs1 signaling and Lkb1-Ampk-Acc signaling was largely recapitulated. Consistent with the drastic metabolic reprogramming following Ar activation, all the metabolic parameters assayed were significantly altered in *Ar*-overexpressing mice exposed to 10% glucose, with no exception. Furthermore, transgenic mice easily developed insulin resistance, fatty liver and obesity. Conversely, Ar deficiency in yellow obese syndrome mice greatly improved the Agouti *A^y^*mutation induced metabolic syndrome. Our results reveal a profound Warburg effect-like metabolic reprogramming as a consequence of abnormal activation of PP/the pseudo-glycolysis. These findings also highlight the important contribution of PP or the putative pseudo-glycolysis in hepatic metabolic homeostasis and the development of metabolic diseases.

## RESULTS

### Glucose-induced Ar activation significantly stimulates the glycolysis and putative pseudo-glycolysis, lactate secretion, oxygen consumption and mitochondrial respiration

We created stable *Ar-*overexpressing AML12 cells (*Ar*-AML12) from normal mouse hepatocyte AML12 cells (NC-AML12) (Fig. 1*A*). No difference in 2-deoxy-glucose uptake was found between the *Ar-*overexpressing AML12 cells and the controls (Fig. 1*B*), suggesting *Ar* overexpression probably will not affect glucose uptake in the liver cells. To investigate the effects of hepatic *Ar* activation on cellular metabolism, we conducted the Seahorse glycolysis and mitochondrial respiration assays with the *Ar*-AML12 and NC-AML12 cells in the presence of high glucose. Upon the injection of 25 mM glucose, the extracellular acidification rate (ECAR) in *Ar*-AML12 cells increased much faster than that of NC-AML12 cells (Fig. 1*C*). Immediately before the addition of oligomycin, ECAR of *Ar-*AML12 cells was already about 43% higher than that of NC-AML12 cells (87.22 ± 5.50 versus 60.75 ± 8.40 mpH/min, *p* < 0.001). Similarly, the glycolytic capacity and glycolytic reserve of *Ar*-AML12 were both much larger than that of NC-AML12 cells. Consistent with the trend of glycolysis, the maximal mitochondrial respiration (oxygen consumption rate, OCR) and spare capacity of respiration were greatly increased in *Ar*-AML12 cells, as compared with that of NC-AML12 cells (Fig. 1*D*). Immediately before the addition of Rotenone/Antimycin A, the maximal oxygen consumption rate for *Ar*-AML12 cells was about 36% higher than that of NC-AML12 cells (405.25 ± 28.82 versus 297.78 ± 36.07 pmol/min, *p* < 0.001). To further verify the increased acidification, we use a chemical kit to assay lactate secretion in *Ar*-AML12 and NC-AML12 cells. Grown in DMEM/F-12 with 17.5 mM glucose for 4 h, the concentration of lactate in the media of *Ar*-AML12 cells was already much higher than that in the media of NC-AML12 cells, which sustained for at least 8 h (Fig. 1*E*). In *Ar*-overexpressing AML12 cells treated with 100 µM Ar inhibitor (ARI), high glucose-induced ECAR and OCR and lactate secretion appeared to be improved. To validate the increased mitochondrial respiration in *Ar*-AML12 cells, we utilized MitoSOX Red to stain the AML12 cells. 4 h after the exposure of 25 mM glucose, a larger amount of superoxide was detected in *Ar*-AML12 cells but not with NC-AML12 cells (Fig. 1*F*). Together these data indicate that in the presence of high glucose, *Ar* overexpression greatly enhances the glycolysis and pseudo-glycolysis and mitochondrial respiration.

**Fig. 1.**
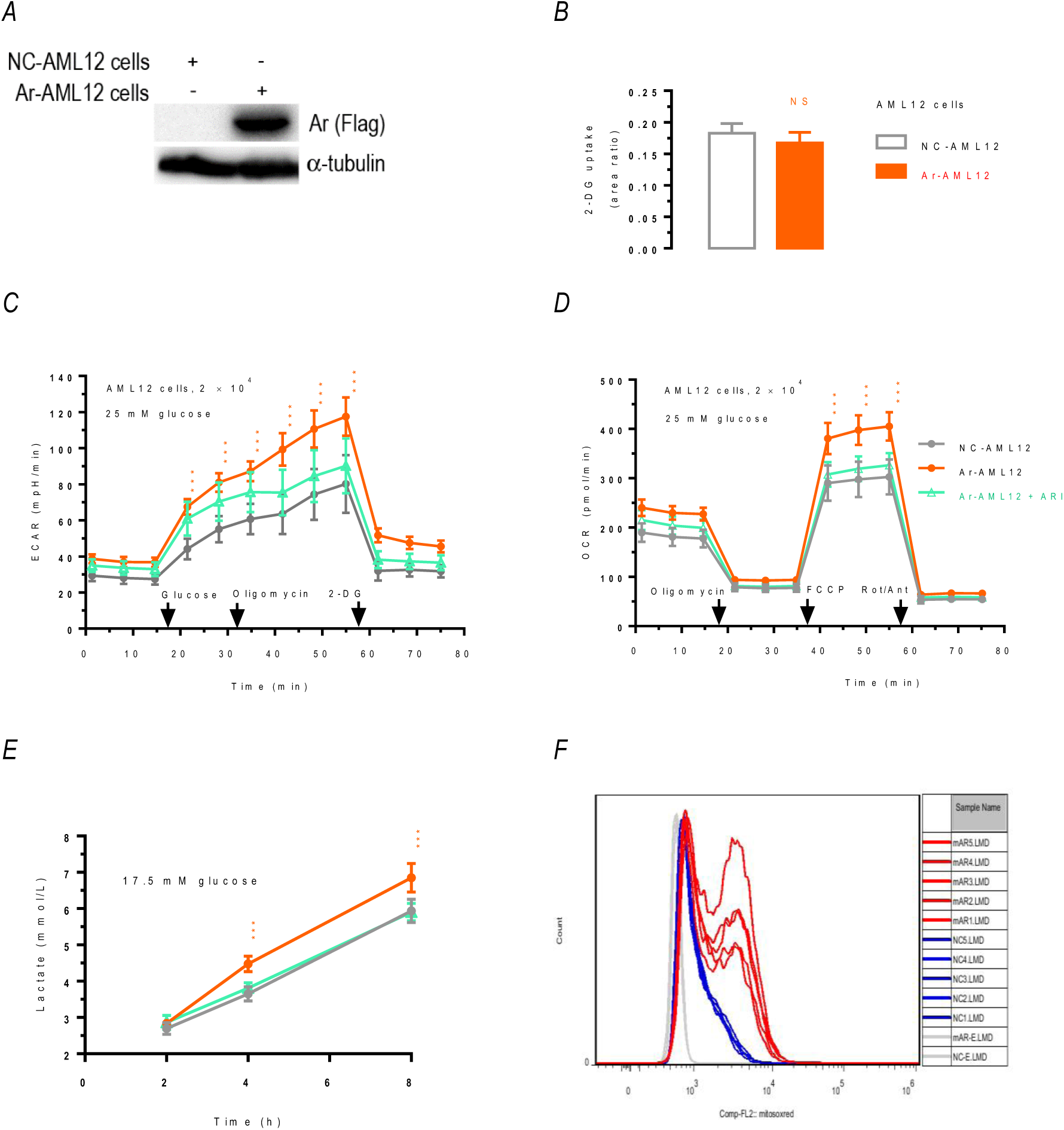
Extracellular acidification, oxygen consumption and mitochondrial superoxide production in Ar-overexpressing AML12 cell and the controls (n = 3). Values were expressed as the mean ± SD. *, *p* < 0.05; **, *p* < 0.01; ***, *p* < 0.001. *A*. Western blots showing Ar overexpression in *Ar*-overexpressing AML12 cells (Ar-AML12) and the control cells (NC-AML12) (*n* = 3). Data were typical for at least three repeated experiments. *B*. 2-deoxy-glucose (2-DG) uptake in Ar-AML12 and NC-AML12 cells. *C.* ECAR in NC-AML12 and Ar-AML12 cells (*n* = 3). Data were typical for at least three repeated experiments. *D*. OCR in NC-AML12 and Ar-AML12 cells. Data were typical for at least three repeated experiments (*n* = 3)2-. *E*. Lactate secretion in NC-AML12 and Ar-AML12 cells (*n* = 3). *F*. Mitochondrial superoxide levels in NC-AML12 and Ar-AML12 cells as assayed by MitoSOX Red (*n* = 2-5). mAR1-5 (red), Ar-AML12 cells; NC1-5 (blue), NC-AML12 cells; NC-E and mAR-E (grey), no dye controls.

### Ar activation triggers Warburg effect -like metabolic reprogramming in the hepatocytes

In order to delineate the metabolomic profile of *Ar*-overexpressing AML12 cells, we performed metabolomic studies through liquid chromatography coupled with mass spectrometry (LC-MS/MS), using cultured NC-AML12 and *Ar*-AML12 cells. In AML12 cells exposed to 25 mM glucose for 4 h, a total of 280 non-redundant metabolites were detected by our LC-MS. Among which, 52 metabolites were significantly altered following Ar activation, with 42 metabolites increased and 10 metabolites decreased by at least 1.5-fold (***p*** < 0.05). Principal Component Analysis (PCA) showed that the PC1 and PC2 scores were 77.3 and 13.0% respectively (Fig. S2***A***), indicating significant differences between NC-AML12 and *Ar*-AML12 cells. Partial Least Squares Discriminant Analysis (PLS-DA) showed increased galactitol, ATP, lactate, urate, NAD^+^ and decreased citrate were among the Variable Importance in Projection (VIP) top scored metabolites (Fig. S2***C***). Enrichment Analysis with 52 significantly changed metabolites (fold change threshold 1.5, ***p*** < 0.05) showed that these metabolites were significantly enriched in metabolic pathways including pyrimidine and purine metabolism, thiamin metabolism, Warburg effect, glycolysis/citric acid cycle, electron transport chain, galactose metabolism, DNL, glycerolipid metabolism *etc.* (Fig. 2***A-B***, Fig. S3). As shown in Fig. 2***A***, the 25 most significantly enriched metabolic pathways were mostly involved in carbohydrate and lipid metabolism, whereas “Lysine degradation” and “Urea cycle” were the only two pathways associated with amino acid metabolism. Remarkably, out of the 58 characteristic Warburg effect associated metabolites, 9 (15.5%) were significantly altered following Ar activation, which included decreased acetyl-CoA and increased GTP, ATP, NADH, D-ribose phosphate, DHAP, thiamine monophosphate, 3-phosphoglycerate and oxoglutaric acid (α-keto-glutaric acid) (Fig. 2***B***, Fig. S4). Meanwhile, the Pathway Analyses indicated that purine/pyrimidine metabolism, thiamine/riboflavin/glutathione metabolism, glycolysis/TCA/pentose phosphate pathway (PPP) and fatty acid elongation and arginine biosynthesis appeared to have the most significant pathway impact scores (Fig. 2***C***). Consistent with the activation of Ar/PP, fructose and galactitol were significantly increased (Fig. S4***A-B***). Ar is known to be able to convert galactose into galactitol (40). Further, lactate, glycerol 3-phosphate, DHAP, ATP, GTP, NADH/NAD^+^ were significantly increased whereas acetyl-CoA was significantly decreased (Fig. S3 & S4).

**Fig. 2.**
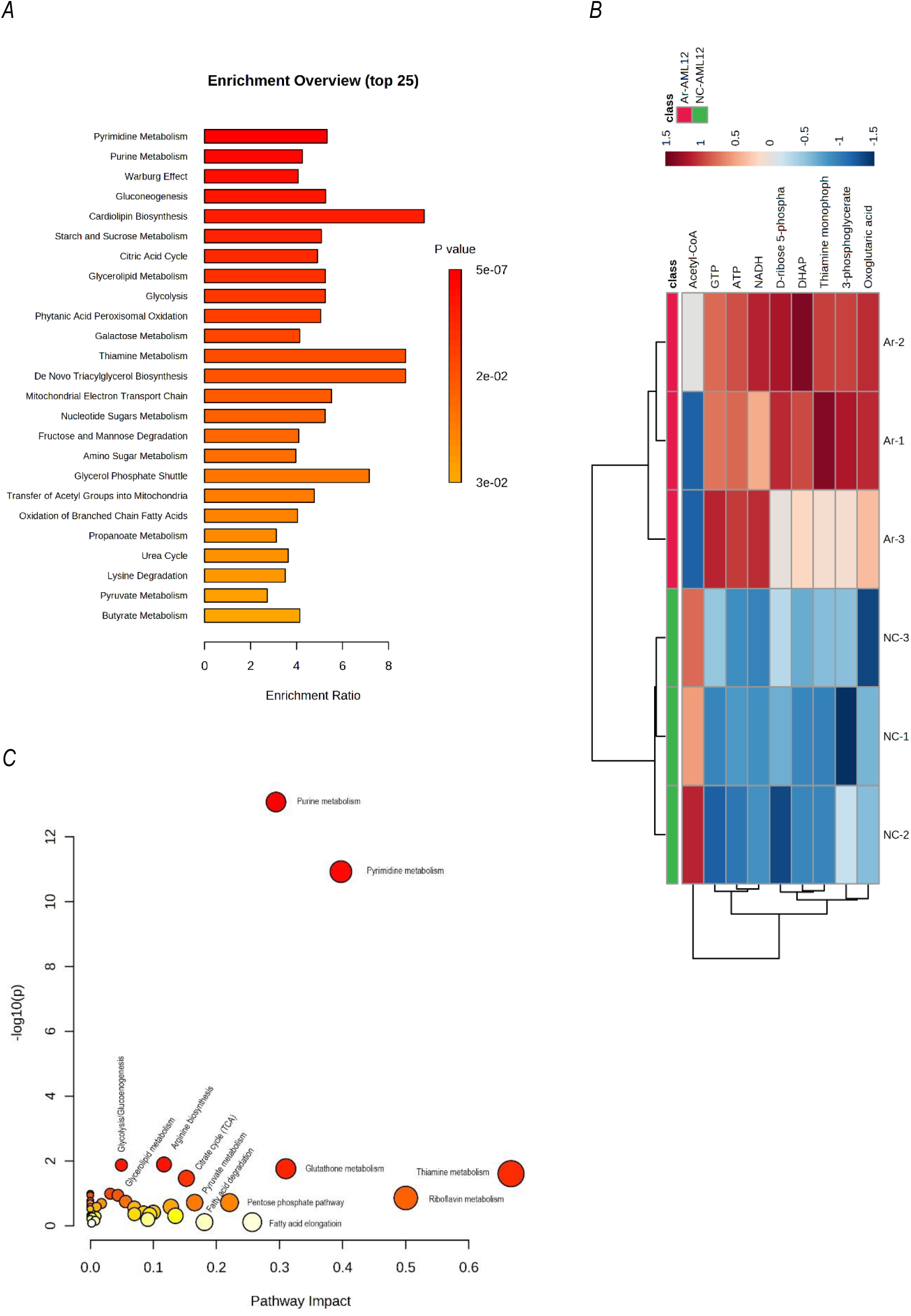
Metabolomic analyses of the control and stable Ar-overexpressing AML12 cells. The LC-MS metabolomic data were analyzed by MetaboAnalyst 5.0 (*n* = 3). ***A***. Metabolic pathway enrichment as a consequence of Ar activation. 52 metabolites that were significantly increased or decreased by a fold change threshold of 1.5 (*t*-test, ***p*** < 0.05) were used for the pathway enrichment analyses. The enrichment ratio is calculated as the number of hits within a particular metabolic pathway divided by the expected number of hits. ***B***. The abundancy of Warburg effect-associated metabolites following Ar activation (*n* = 3). The heap map was generated with 9 metabolites that were significantly increased or decreased by a fold change threshold of 1.5 (*t*-test, ***p*** < 0.05). NC-1∼3, NC-AML12 cells; Ar-1∼3, Ar-AML12 cells. ***C***. Pathway Impact scores based on 52 significantly changed metabolites (fold change threshold 1.5, *t*-test, ***p*** < 0.05).

Although NADP^+^/NADPH was not found to be significantly altered as analyzed by LC-MS (Fig. S4***U***), a separate chemical kit analyses confirmed the increase of NADP^+^/NADPH following Ar activation (Fig. S2***B***). Surprisingly, an unusual by-product of PPP and the conventional glycolysis, sedoheptulose 1,7-bisphosphate (SBP), was significantly increased in *Ar*-overexpressing cells (Fig. S4***V***). SBP is known to be involved in the regulation of PPP (41) and a previous finding suggests that the level of SBP in hepatoma cells was greatly increased by ROS exposure (42). Ar-mediated increase in SBP thus probably in part reflects Ar-mediated enhancement of mitochondrial respiration and ROS production (Fig. 1***B*** & 1***C***). Also unexpectedly, both reduced glutathione (GSH) and oxidized glutathione (GSSG) were not decreased following Ar activation, they actually increased slightly but not significantly (Fig. S3***D*** & S4***AA***). These metabolomic data thus indicate that Ar activation drastically impact major metabolic pathways encompassing the metabolism of carbohydrates, lipids, and purine or pyrimidine bases, causing a drastic and profound Warburg effect-like metabolic reprogramming in the liver cells.

### Irs1 and Lkb-Ampk signaling is significantly altered by Ar overexpression or knockdown

Insulin receptor substrate-1 (Irs1) is a protein that plays a key role in insulin signaling and deficiency in Irs1 in adipocyte is known to predict insulin resistance and type II diabetes (43). On the other hand, AMP-activated protein kinase (Ampk) is a master regulator that is involved in the regulation of carbohydrate and lipid metabolism, mitochondrial and lysosomal homeostasis and DNA repair, thereby contributing to the development of cancer, obesity, diabetes, nonalcoholic steatohepatitis and other disorders (44). Ampk is activated by its upstream activator liver kinase B1 (Lkb1) through phosphorylating Ampkα at threonine-172. Further, Ampk inhibits fatty acid and cholesterol biosynthesis and promote fatty acid oxidation in part through phosphorylating acetyl-CoA carboxylase (Acc) at serine-79 to suppress its activity (45). To explore Ar/PP-mediated alterations in metabolic signaling, we transiently overexpressed or knocked down *Ar* in AML12 cells (Fig. S5***A***). Following plasmid transfection for 24 h or lentiviral transfection for 96 h, *Ar* overexpression significantly reduced whereas *Ar* knockdown increased mRNA (Fig. S5***B***) and protein expression of Irs1 (Fig. 3***A***). On the other hand, despite that both *Ar* overexpression and *Ar* knockdown did not appear to significantly change the protein expression of Lkb1, Ampkα and Acc, the phosphorylated proteins (pLkb1^S428^, pAmpkα^T172^ and pAcc^S79^) were reduced in *Ar-*overexpressing cells but increased in *Ar*-knockdown cells, resulting in significant changes in the ratios of pLkb1^S428^/Lkb1, pAmpkα/Ampkα and pAcc^S79^/Acc (Fig. 3***A***). In AML12 cells grown in 25 mM glucose, Ar protein expression was time dependently increased (Fig. 3***B***). With the increased expression of Ar protein, Irs1 steadily increased in the first 6 h but thereafter slowly reduced up to 48 h. Similar trend of protein expression was observed for the phosphorylated proteins pLkb1^S428^, pAmpkα^T172^ and pAcc^S79^ but not the unphosphorylated. In contrast to 25 mM glucose exposure, the effects of 5 mM fructose were much less pronounced (Fig. 3***C***). Overall, these data indicate that Ar negatively regulates Irs1 and Lkb1-Ampk signaling and this metabolic signaling should be part of the Ar/PP-mediated metabolic reprogramming in the hepatocytes.

**Fig. 3.**
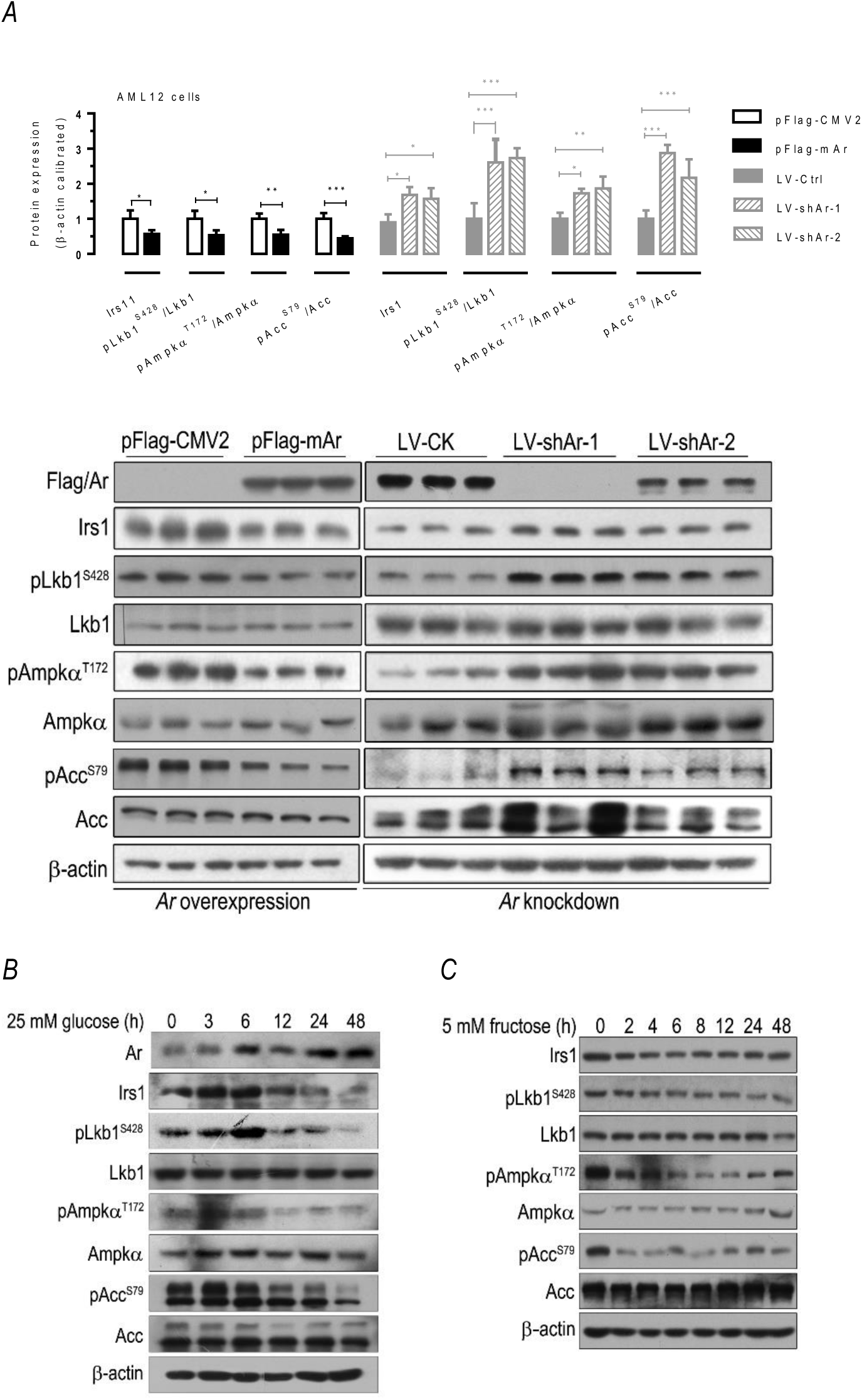
Irs-1 and Lkb1-Ampk signaling in Ar overexpressing or knockdown AML12 cells. ***A.*** Overexpression of *Ar* down-regulated whereas lentivirus-mediated *Ar* knockdown up-regulated protein expression/phosphorylation of Irs-1, Lkb1, Ampkα and Acc. AML12 cells were transiently transfected with pFlag-CMV2 or pFlag-mAr or infected with lentiviruses pLV-CK, pLV-shAr-1 or pLV-shAr-2 respectively. 24 h after the transfection or 96 h after the infection, cells were collected for Western blots. Experiments were performed with at least three separate samples (*n* = 3) analyzed in triplicate. Values were expressed as the mean ± SEM. NS, not significant; *, ***p*** < 0.05; **, ***p*** < 0.01; ***, ***p*** < 0.001. ***B***. Effects of 25 mM glucose exposure on protein expression/phosphorylation of Ar, Lkb1, Ampkα, Acc and Irs-1. AML12 cells were cultured in the presence of 25 mM glucose. Cells were collected at 0, 3, 6, 12, 24 and 48 h respectively. ***C***. Effects of 5 mM fructose exposure on protein expression/phosphorylation of Lkb1, Ampkα, Acc and Irs-1. AML12 cells were cultured in the presence of 5 mM fructose. Cells were collected at 0, 2, 4, 6, 8, 12, 24 and 48 h respectively.

### Ar mediates DNL and TG accumulation

Since at least 4 out of 9 putative DNL associated metabolites were enriched following Ar activation (Fig. S3***C***) and *Ar* knockdown increased the phosphorylation of the key fatty acid synthesis enzyme Acc at serine-79 by Ampk to suppress its lipogenesis activity (Fig. 3***A***), we wanted to verify the effects of Ar overexpression or inhibition on lipid accumulation and DNL in hepatocytes. 24 h after the exposure of 25 mM glucose, *Ar*-overexpressing AML12 (*Ar*-AML12) cells showed more significant formation of lipid droplets than normal control AML12 (NC-AML12) cells, whereas inhibition of Ar with ARI significantly reduced the formation of lipid droplets (Fig. 4***A***). This result verified the effect of Ar on hepatocyte lipid accumulation.

**Fig. 4.**
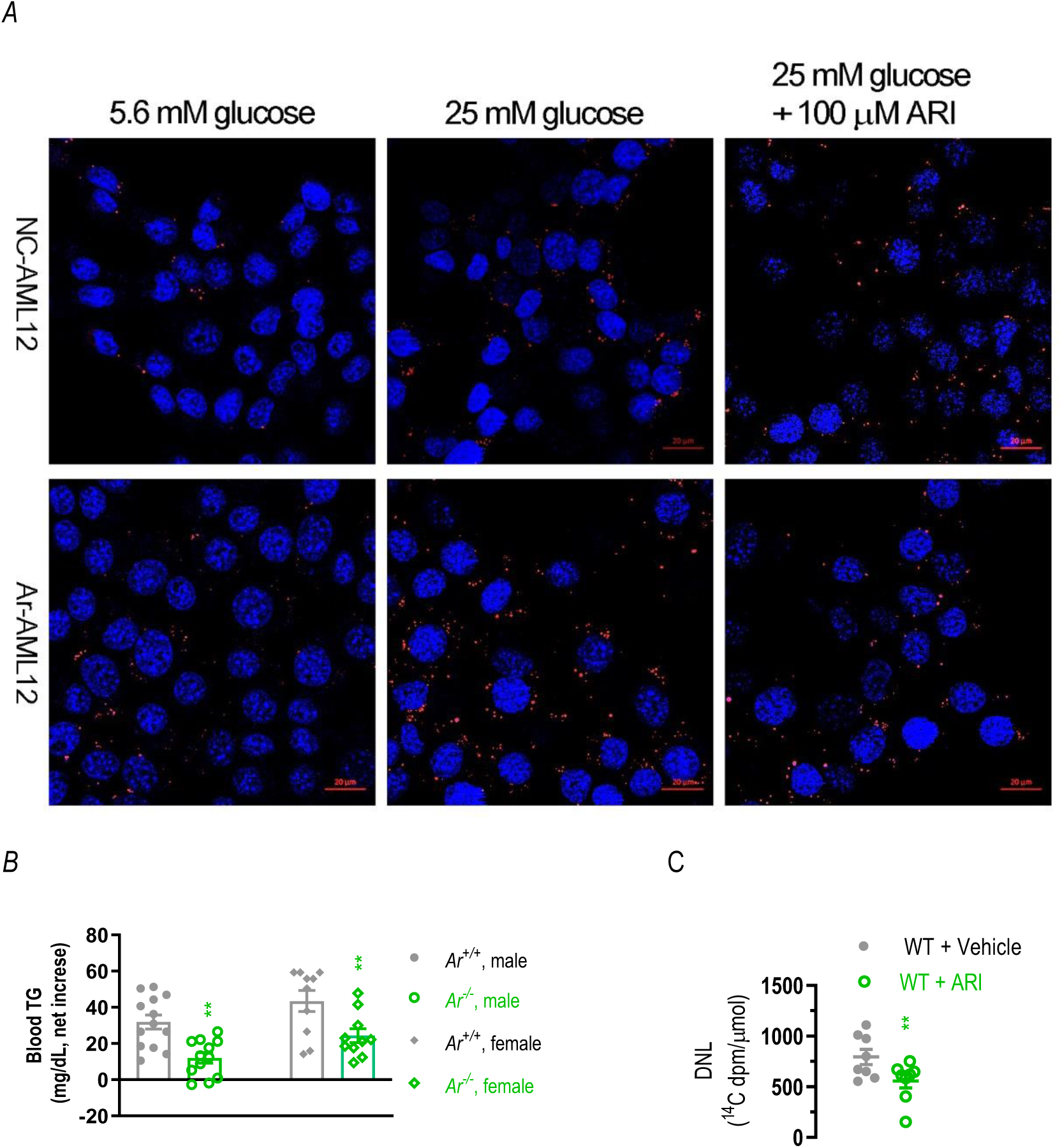
High glucose-induced lipid synthesis. ***A.*** High glucose-induced formation of lipid droplets in AML12 cells. NC-AML12 and Ar-AML12 cells were cultured in DMEM containing 5.6 mM glucose for 14 h. Cells were then cultured in media containing either 25 mM glucose or 25 mM glucose + 100 µM ARI for another 24 h, respectively. Then cells were fixed in 3% paraformaldehyde fix solution, stained with Nile red (0.05 μg/ml) (Sigma, Cat # N3013) for 10 min, then washed three times with PBS before being visualized by confocal laser scanning microscopy. Results were typical of at least three experiments. ***B***. Blood levels of TG in *Ar^+/+^*and *Ar^-/-^* C57BL/6 mice following a peritoneal injection of glucose (4 g/kg body weight, *n* = 10-13). **, *P* < 0.01. ***C***. Hepatic DNL (^14^C-U-glucose incorporation into hepatic TG). wild type C57BL/6 mice (WT) with or without pretreated ARI pretreatment were injected ^14^C-U-glucose as described. 1 h after radio-active ^14^C glucose loading, liver tissues were dissected and lipid extracted for separation by thin layer chromatography. Lipid spots were identified and collected for scintillation counting (*n* = 8). *, ***p*** < 0.05 between ARI-treated and untreated C57BL/6 mice.

To demonstrate the *in vivo* effects of Ar-mediated DNL, we intraperitoneally injected a bolus dose of 4 g/kg body weight into the wildtype control and *Ar*-null C57BL/6 mice. 1 h after the glucose loading, glucose-induced increase in blood TG levels in both male and female *Ar*-null C57BL/6 mice were significantly lower than that in the control mice (Fig. 4***B***). For example, the average net increase of blood TG in male *Ar*-null C57BL/6 mice was only about 37.9% of that in the male control mice (31.93 ± 3.923 versus 12.1 ± 2.89, ***p*** < 0.01). To further demonstrate DNL in the liver, we intraperitoneally injected into wildtype C57BL/6 mice and wild type C57BL/6 mice pretreated with Ar inhibitor zopolrestat (ARI, 50 mg/kg body weight) with a glucose solution (2 g/kg body weight) containing radioactive ^14^C-U-glucose (40 Ci of ^14^C-U-glucose/mouse). 1 h after radio-active ^14^C glucose loading, the liver tissues were dissected and lipid extracted for separation by thin layer chromatography. Lipid spots were identified and collected for scintillation counting. The results showed that significantly less ^14^C-labeled glucose was converted into newly-synthesized (radio-labeled) TG in the liver of ARI-pretreated livers than that of the vehicle-treated mice (556.3 ± 67.26 versus 795.2 ± 75.27 ^14^C dpm/μmol TG, ***p*** < 0.05, Fig. 4***C***). These data verify that Ar activation is tightly linked with lipid accumulation and DNL in the liver cells.

### Sustained AR activation in transgenic mice leads to the development of insulin resistance, fatty liver and obesity

To link the cellular findings from *Ar*-overexpressing AML12 cells mentioned above to *in vivo* mouse models, we first created lines of transgenic mice (Tg1 and Tg2) to stably overexpress human *AR* liver-specifically (Fig. S6***A***-***D***). In 154-d old Tg1 and Tg2 mice received 10% glucose in drinking water from the weaning date, liver-specific AR activation again appeared to cause a small but not significant reduction in protein expression of Irs1, and phosphorylated Irs1, Lkb1, Ampkα and Acc but not the unphosphorylated protein (Fig. S6***E***-***F***), which further corroborated the regulation of Irs1 and Lkb1-Ampk signaling by AR. Regardless of high glucose feeding or regular chow feeding only, *AR*-overexpressing transgenic mice (Tg1) gained weight more quickly than the wildtype controls, although 10% glucose supplementation further accelerated the weight gain (Fig. 5***A***). By the age of 154-d, Tg1 on 10% glucose developed significant fatty liver (Fig. 5***B***). Meanwhile, hyperinsulinemia and hyperleptinemia and insulin resistance were observed for both Tg1 and Tg2 on 10% glucose (Fig. 5***C-E***). Other metabolic parameters including the serum levels of alanine aminotransferase (ALT), fructose, TG, cholesterol and the liver weight, the liver levels of fructose, TG and cholesterol were all significantly increased for both Tg1 and Tg2 (Fig. 5***F-M***). These *in vivo* data are consistent with the drastic hepatic metabolic reprogramming following Ar activation. Sustained livre-specific activation of AR/PP, therefore, appears to be sufficient to cause metabolic alterations leading to the development of insulin resistance, fatty liver, obesity and other abnormalities.

**Fig. 5.**
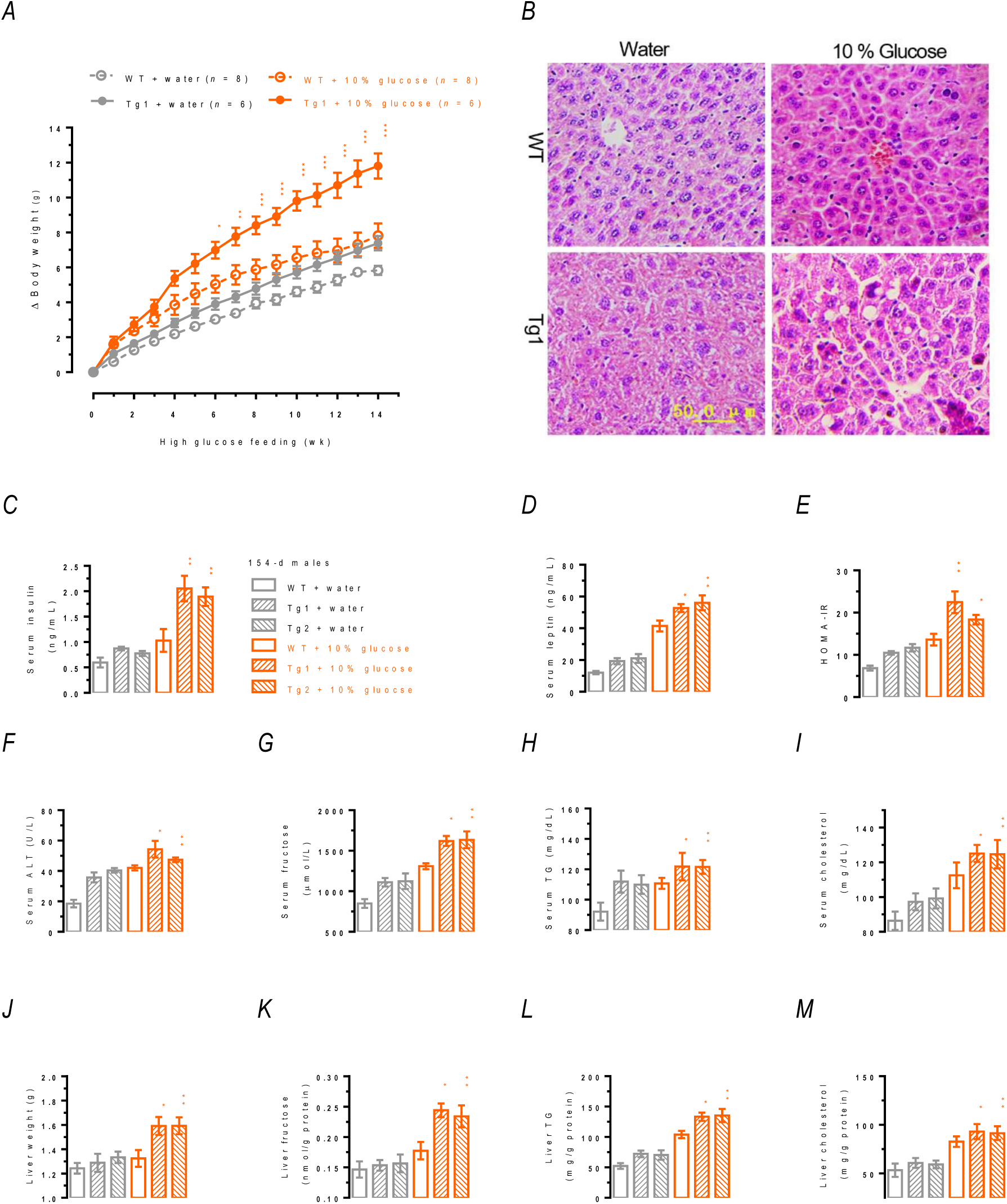
Effects of Liver-specific Ar transgenic overexpression on obesity, liver steatosis and other metabolic parameters. Values were expressed as the mean ± SEM. Pair-wise comparisons were made for WT + glucose versus Tg1 + glucose (in yellow *). Only male mice at the age of 154-d were used. *, *P* < 0.05; **, *P* < 0.01; ***, *P* < 0.001. ***A***. Body weight gain (*n* = 6-8). 10% glucose drinking water was started from the day of weaning. ***B***. Liver steatosis by H&E staining (*n* = 3). Magnification, 200×. ***C-M***. Serum insulin (***C***), HOMA-IR (***D***), serum ALT (***E***), leptin (***F***), serum fructose (***G***), serum TG (***H***), serum cholesterol (***I***), liver weight (***J***), liver fructose (***K***), liver TG (***L***) and liver cholesterol (***M***) (*n* >= 3).

### Suppression of Ar-mediated metabolic reprogramming greatly improves Agouti A^y^ mutation-induced insulin resistance, fatty liver and obesity

Agouti and Agouti-related proteins are important signal proteins involved in the regulation of the melanocortin receptor signaling (46). Mice carrying the dominant *Agouti* mutant allele *A^y^* develop the so-called yellow obese syndrome (46,47). The yellow obese syndrome in mice encompasses many pleiotropic effects including hyperphagia, yellow fur, hyperglycemia, hyperinsulinemia, hyperleptinemia, insulin resistance, obesity, type II diabetes and increased susceptibility to neoplasia. The Agouti yellow obese mice (C57BL/6, *A^y^/a*) therefor represent one of the more physiologically-relevant mouse models for metabolic studies. To evaluate the feasibility of treating or preventing the metabolic syndrome including insulin resistance, fatty liver and obesity through suppression of *Ar*-mediated metabolic reprograming, we crossed the Agouti yellow mice with *Ar*-knockout C57BL/6 mice (Fig. S7*A*), generating the Agouti yellow obese mice carrying *Ar*-null mutation and its controls, *i.e.*, two groups of mice with non-yellow fur color (*a/a::Ar^+/+^* and *a/a::Ar^-/-^*) and two groups of mice with yellow color fur (*A^y^/a::Ar^+/+^ and A^y^/a::Ar^-/-^*). In comparison with the 130-d old control mice (*a/a::Ar^+/+^* and *A^y^/a::Ar^+/+^* ), the liver protein expression (Fig. 6*A*) of Irs1 and the phosphorylated proteins of Lkb1, Ampkα and Acc were in general increased in the two groups of *Ar*-deficient mice (*a/a::Ar^-/-^* and *A^y^/a::Ar^-/-^*) under normal feeding conditions (regular chow and water *ad libitum*), except Acc and pAcc^S79^ in the *A^y^/a::Ar^-/-^* mice. Consistent with the yellow obese syndrome, blood glucose, liver fructose, and serum levels of fructose, insulin, leptin and TG were all significantly increased in the 130-d old yellow mice with normal *Ar* (Fig. S7*B-G*). *Ar* deficiency in the yellow mice, however, at least resulted in improvement in blood glucose and serum fructose. As compared to the yellow mice with normal *Ar* (*A^y^/a::Ar^+/+^*), however, yellow mice lacking *Ar* (*A^y^/a::Ar^-/-^*) appeared to have greatly improved liver histology (Fig. 6*B* & Fig. S7*H*). In comparison with that of the control yellow mice (*A^y^/a::Ar^+/+^*), obesity and fat tissue growth in *Ar*-deficient (*A^y^/a::Ar^-/-^*) yellow mice were also significantly improved (Fig. 6*C-D*), with no significant difference in chow consumption (Fig. S7*I*). Results from glucose tolerance tests, insulin sensitivity and HOMA-IR tests (Fig. 6*E-G*) suggested that *Ar* deficiency greatly improved glucose intolerance and insulin insensitivity in the yellow mice. Collectively these data showed that most of the abnormal phenotypes of the yellow obese syndrome, which include hyperglycemia, insulin resistance, fatty liver and obesity, were greatly improved in mice lacking *Ar. Ar* deficiency probably achieved this at least in part through upregulating Irs1 and Lkb1-Ampk signaling. Importantly, our data suggest the feasibility and efficacy of treating or preventing insulin resistance, fatty liver, obesity and other metabolic disorders through suppression of hepatic Ar/PP.

**Fig. 6.**
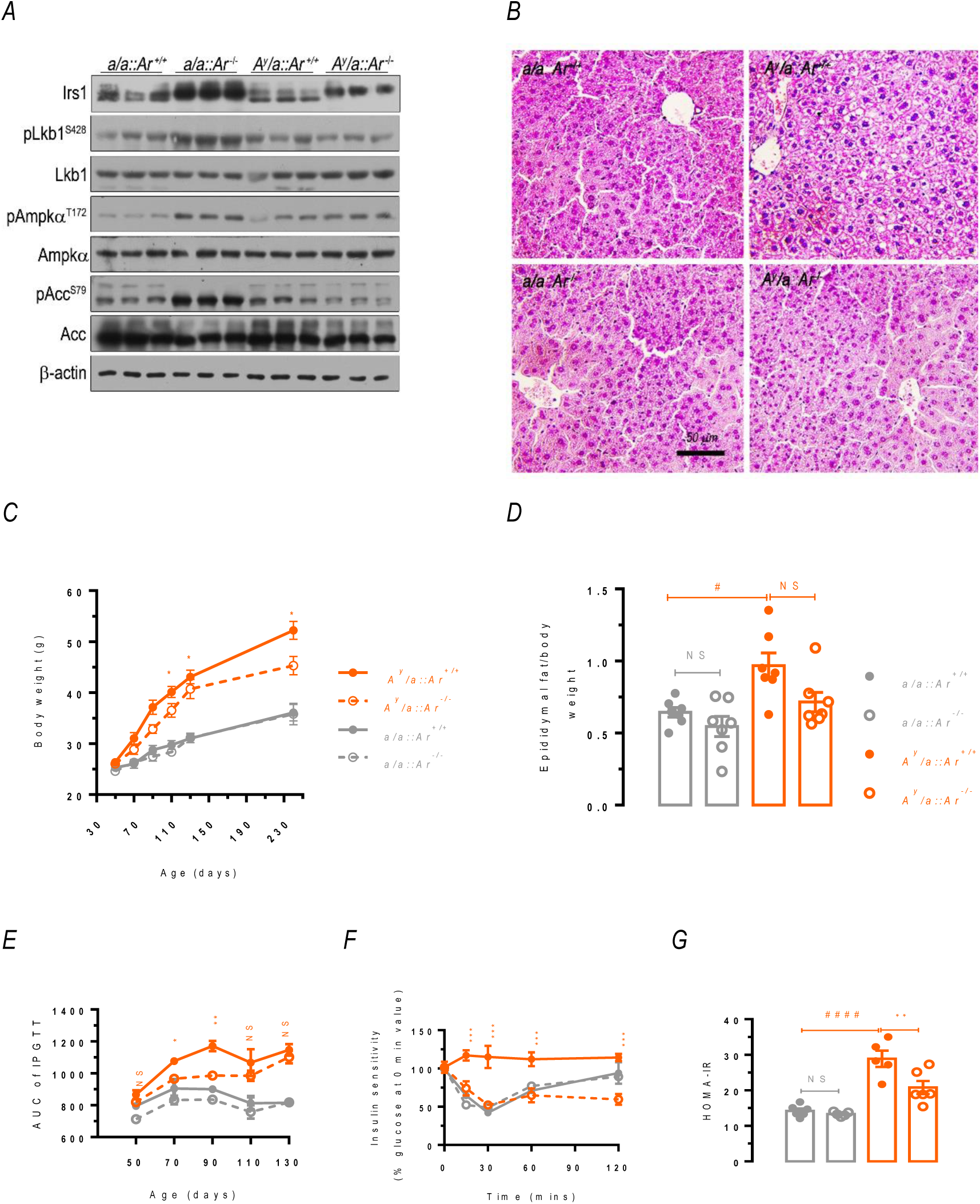
Effects of Ar deficiency on protein expression and phenotypes of Agouti mice and the controls in 4 groups of mice at the age of 130-d. Values are expressed as the mean ± SEM. Pair-wise comparisons were made for a/a::Ar^+/+^ versus a/a:: Ar^-/-^ (in grey *) or a/a::Ar^+/+^ versus A^y^/a::Ar^+/+^ (in yellow #) or A^y^/a::Ar^+/+^ versus A^y^/a::Ar^+/+^ (in yellow *) . NS, not significant; * or #, p < 0.05; **, p < 0.01; ####, p < 0.0001. Only male mice were used. A. Hepatic protein expression/phosphorylation of Irs-1, Lkb1, Ampkα and Acc (n = 3). B. Liver steatosis (n >= 3). Magnification, 200×. C. Time dependent body weight gain (n = 7-16). D. Weights of epididymal fat pads (n = 7). E. Glucose tolerance tests (n = 5). AUC, area under curve. F. Insulin sensitivity tests (n = 3). E. HOMA-IR tests (n = 5-6).

## DISCUSSION

Hepatic expression of Ar in rodents and humans are known to be low or undetectable under normal physiological conditions (48,49). Upon high glucose exposure, however, Ar can be significantly upregulated in AML12 cells and the liver of diabetic mice (30). In human Chang liver cells, proteome analyses show that Ar is the most significantly up-regulated protein following exposure of 25 mM glucose (50). In humans and rodents, greatly increased hepatic Ar expression is often associated with the development of liver diseases including HBV/HCV, alcoholic liver disease, autoimmune hepatitis, and HCC (38,48,49,51–54). Hepatic Ar expression therefore, is highly inducible. In contrast to Ar, other Ar down-stream enzymes unique for the putative pseudo-glycolysis, which include Sdh, Khk-A/C, aldoB and Tkfc, are well-equipped in the liver (10). Carbohydrate response element binding protein (Chrebp) is a transcriptional factor playing important roles in regulating glucose and lipid metabolism, especially lipogenesis, glycogen synthesis, and gluconeogenesis. Interestingly, the fructose transporter *Glut5* and fructolytic genes *Khk*, *AldoB*, *Tkfc* are all under the transcriptional control of Chrebp, which can be activated by both glucose and fructose (10). In another study, Tkfc was found to play a gatekeeper role through coupling fructolysis with lipogenesis and fructose tolerance (6). The putative pseudo-glycolysis in the liver therefore not only is lipogenic but also operates more efficiently than the conventional glycolysis, since it lacks the tight feedback regulation as seen in the conventional glycolysis.

The drastic metabolic reprogramming brought about by Ar activation is of particular importance to the elucidation of hepatic glucose and lipid homeostasis. As expected, activation of Ar/PP/the putative pseudo-glycolysis in liver cells leads to significant accumulation of fructose, galactitol and lactate, together with significant increase in ATP and NADH and the depletion of NADPH. Moreover, hepatic Ar activation was shown to suppress whereas Ar inhibition was shown to enhance Irs1 signaling and Lkb-Ampk-Acc signaling. Further, Ar/PP activation promotes DNL. Previous studies indicate that fructose and galactose are both lipogenic (7,55). A recent study further suggests that in addition to glucose and fructose, galactose but not mannose, L-arabinose, xylose and ribose enhances hepatic fat accumulation in HepG2 cells (56). Manifested by increased galactitol following Ar activation, enhanced galactose metabolism (Fig. S3***B***) can also be expected to promote DNL.

Surprisingly, the activation of Ar/PP/the putative pseudo-glycolysis strongly impacted the metabolism of pyrimidine/purine and thiamin/riboflavin. How the activation of Ar/PP/the pseudo-glycolysis causes systematic alterations of pyrimidine/purine and thiamin/riboflavin is not clear at the moment. However, most of these pyrimidine/purine derivatives, thiamin and riboflavin are either the co-enzymes/co-factors or their precursors deeply involved metabolic reactions and mitochondrial function. As glycolysis/PPP is greatly enhanced following Ar/the putative pseudoglycolysis activation, it is conceivable that these co-enzymes/co-factors (e.g., thiamin (vitamin B1)/riboflavin (vitamin B2), ATP/GTP, NAD^+^/NADH, NADP^+^/NAPDH etc.) will be in large demand. Moreover, when Tkfc is activated, its cyclizing lyase activity will split FAD to AMP and riboflavin cyclic-4,5-phosphate (cyclic FMN or cFMN) to contribute to enhanced pyrimidine/purine metabolism (10,25).

While we demonstrate that increased glucose flux through PP/the putative pseudo-glycolysis leads to significantly increased OCR and mitochondrial ROS production (Fig. 1***D******&***F), both reduced glutathione (GSH) and oxidized glutathione (GSSG) were not significantly reduced (Fig. S3***B***, Fig. S4***AA***). This is in conflict with a long-held view that overactivation of PP will cause oxidative stress by depleting NADPH to deplete GSH (29,57). On the other hand, increased glucose flux through PP leads to significantly increased “glycolysis” as indicated by increased ECAR in the glycolysis analyses (Fig.1***C***). This increase in “glycolysis”, however, represents more of the putative pseudo-glycolysis rather than the conventional glycolysis, as this increase is caused mainly by Ar activation.

In liver-specific *Ar*-overexpressing transgenic mice exposed to 10% glucose drinking water, Ar-mediated metabolic reprogramming appeared to have been translated into altered metabolic parameters and phenotypes consistent with the metabolic alterations, which include hyperinsulinemia, hyperleptinemia, hypertriglyceridemia, hypercholesterolemia, insulin resistance, fatty liver and obesity. In addition, Ar activation-induced suppression of Irs1 signaling and Lkb-Ampk-Acc signaling as seen in the AML12 cells was largely recapitulated in *Ar*-overexpressing transgenics exposed to 10% glucose drinking water. Suppression of Irs1 in the *Ar*-overexpressing transgenics at least in part explains the development of insulin resistance (43,58,59). We previously have shown that Ar overexpression suppresses fatty acid oxidation through regulating the phosphorylation and activity of peroxisome proliferator activated receptor α (PPARα) (30). In this investigation, we further demonstrate that Ar/PP activation in both *Ar*-overexpressing AML12 hepatocytes and liver-specific *Ar*-overexpressing transgenics suppresses Lkb-Ampk-Acc signaling, leading to the release of fatty acid synthesis activity of Acc to promote DNL (44,45). Ar/PP activation, therefore, promotes fatty acid synthesis while suppresses fatty acid oxidation, the net results will therefore be the accumulation of TG and fat. In the yellow obese mice, the blockage of Ar/PP significantly ameliorated yellow obese mice-associated phenotypes of insulin resistance, fatty liver and obesity. These findings provide important mechanistic insights with regard to how aberrant activation of Ar/PP/the putative pseudo-glycolysis might lead to the development of relevant metabolic diseases.

Otto Warburg showed 100 years ago that, under aerobic conditions, cancer tissues metabolized approximately tenfold more glucose to lactate in a given time than normal tissues, a phenomenon now known as the Warburg effect (60). The Warburg effect holds that cancer cells preferentially utilize glycolysis for energy production even in the presence of oxygen (aerobic glycolysis). While the exact mechanisms are still being investigated, it is believed to be driven in part by oncogenic signaling pathways that alter the expression of key metabolic enzymes and transporters (61). Although the Warburg effect was initially discovered in the cancer cells, recent studies suggest that the Warburg effect might also be involved in non-tumor disorders including pulmonary hypertension, neuronal disorders, cardiovascular diseases, and kidney diseases (62,63).

Most remarkably, we showed that following the activation of Ar/PP/the pseudo-glycolysis, AML12 cell underwent a drastic Warburg effect like metabolic reprogramming. And this appeared to happen in the presence of enhanced rather than defective mitochondrial respiration. Indeed, 9 out of 58 Warburg effect-associated metabolites were significantly altered. These characteristic metabolites included decreased acetyl-CoA and increased GTP, ATP, NADH, D-ribose phosphate, DHAP, thiamine monophosphate, 3-phosphoglycerate and oxoglutaric acid (Fig. 2***B***, Fig. S4). The reason for the decrease in acetyl CoA is not clear but very likely is due to the enhanced DNL. In addition to the alterations in Warburg effect-associated metabolites, other parameters for Warburg effects, such as ATP and lactate and DNL were also significantly increased. Our data thus suggest strongly that the aerobic glycolysis probably is not the only contributor of the Warburg effect.

Studies have suggested that Ar/PP and fructose could be implicated in cancer cell metabolism, proliferation and aggressiveness (27,37,64–69). To explain why obesity and metabolic syndrome are closely linked with the development of cancers, it has been suggested that both exogenous and endogenous fructose contribute to the Warburg effect in cancer cells through increasing glycolysis and suppressing oxidative phosphorylation, while providing biosynthetic precursors and through production of uric acid and lactic acid. (65). In particular, uric acid is thought to facilitate carcinogenesis by inhibiting the TCA cycle, stimulating cell proliferation by mitochondrial ROS, and blocking fatty acid oxidation. Lactic acid, on the other hand, is thought to contribute to cancer growth by suppressing fat oxidation and inducing oncogene expression. Interestingly, it has recently been demonstrated that in hepatic cancer cells, overexpression of Ar led to indications of the Warburg effect, which included enhanced expression of cMYC, hexokinase II, KHK, lactate dehydrogenase A, and increased lactate secretion (38). Conversely, inhibition of PP resulted in suppression of the metabolic reprogramming. Consistent with results from cancer cells, our current demonstration of the drastic Warburg effect-like metabolic reprogramming in non-cancerous AML12 hepatocytes provides a strong experimental support for the notion that PP or the putative pseudo-glycolysis is an important contributor for the Warburg effect in both cancer cells and other proliferating cells.

In summary, we reveal that in liver cells, the activation of Ar/PP/the putative pseudo-glycolysis will trigger and drive a drastic metabolic reprogramming similar to that of the Warburg effect in cancer cells. Moreover, insulin signaling and Ampk signaling are also significantly altered. In normal subjects, Ar/PP mediated metabolic reprogramming tends to promote DNL, insulin resistance, fatty liver and obesity. In cancer cells, we expect Ar/PP mediated metabolic reprogramming will be part of the Warburg effect to provide necessary energy and metabolic intermediates to support the growth and proliferation of cancer cells. Our results have important implications for hepatic glucose and lipid metabolism and energy homeostasis. Our findings also shed light on the development of novel therapeutic strategies and drugs for diseases including fatty liver, obesity, and cancers, through targeting Ar/PP mediated Warburg effect-like metabolic alterations.

## MATERIALS AND METHODS

### Cell culture and treatments

Mouse AML12 hepatocytes were purchased form the ATCC (Manassas, VA, USA) and were normally cultured in a DMEM/F-12 (GIBCO, Grand Island, NY, USA) medium containing 5% fetal bovine serum (GIBCO). Plasmid DNA transfection was performed with Lipofectamine 2000 reagent (Invitrogen) according to the manufacturer’s instructions. For lentivirus-mediated AR knockdown, AML12 cells were transduced with viruses with the aid of 8 ug/ml polybrene (Sigma) for 72 h.

To prepare AML12 *cell* stably overexpressing *Ar,* AML12 cells were infected with the control lentivirus (pLV-TagII) or *Ar*-overexpressing lentivirus (pLV-TagII-mAR) packaged in HEK293T cells for 24 h. Stable cells were selected with puromycin (2 ug/ml) for 2-4 d, yielding a line of stable *Ar*-overexpressing AML12 cell (*Ar*-AML12) and its control (NC-AML12). *Ar* overexpression in *Ar*-AML12 was confirmed with Western blots using anti-Flag antibody (Fig. S1A). Cells were maintained in DMEM/F-12 media until use.

### Animals

To generate transgenic mice overexpressing human *AR* in mouse liver, *AR* cDNA were first amplified from human LO2 cells and subcloned into pFlag-CMV2. A Flag-AR fragment were PCR-amplified with the primers CAG<u>ATCGAT</u>AT-GGACTACAAAGACGATGAC and ACT<u>CTCGAG</u>ATCCTCTAGAGACGAGCAGGC, which carry a ClaI site and a XhoI site, respectively (underlined) and subcloned into pLIV-Le6, a vector containing the constitutive human *ApoE* gene promoter and its hepatic control region (a gift from Dr. John Taylor, Gladstone Institute, San Francisco, CA) to drive *AR* expression liver-specifically. The NotI-SpeI fragment containing the liver-specific ApoE promoter as well as the human cDNA (Fig. S6A) was then isolated and used for pronuclear microinjection with FVB mouse eggs. The presence of transgene was assayed by PCR of genomic tail DNA and Western blots of transgenic proteins, using antibodies listed on Table S1 and Flag-hAR primers listed in Table S1.

Mice deficient in *Ar* and the genotyping were described previously (70). C57BL/6 mice carrying *Ar^-/-^*mutation and the control mice were then mated with the Agouti yellow mice (C57BL/6 *A^y^/a*, Jackson Laboratory, Bar Harbor, Maine) to obtain 4 groups mice, i.e., *A^y^/a::Ar^+/+^*, *A^y^/a::Ar^-/-^, a/a::Ar^+/+^*, and *a/a*::*Ar^-/-^* respectively. The dominant *A^y^* mutation in the agouti yellow mice causes the development of insulin resistance, fatty liver and obesity, and a readily recognizable yellow coat color (46,71). Yellow coat color indicates a heterozygous genotype of *A^y^/a* (*A^y^* homozygosity is lethal). Yellow mice can therefore be easily distinguished from mice with non-yellow coat colors (*a/a*).

The mice were maintained in animal rooms under specific-pathogen free conditions with a regular 12-hr light and 12-hr darkness cycle. These minimal disease areas were maintained at a temperature of 22 °C and a relative humidity of 50%. The mice were allowed free access to regular chow and tap water unless specified otherwise.

All experimental procedures involving animals were performed in accordance with animal protocols approved by the Institutional Animal Use and Care Committee of Xiamen University

### Glycolysis stress tests and mitochondrial stress tests in AML12 cells

Glycolysis stress and mitochondrial stress were determined by assaying the extracellular acidification rate (ECAR) and oxygen consumption rate (OCR) using the Seahorse Bioscience Extracellular Flux Analyzer (XF96; Seahorse Bioscience) in real time at 37 °C. Three groups of AML12 cells were used, namely, normal AML12 cells (NC-AML12), *Ar*-overexpressing stable AML12 (*Ar*-AML12) cells and *Ar*-AML12 cells supplemented with 100 µM Ar inhibitor zopolrestate (*Ar*-AML12 + ARI). Briefly, 2 × 10^4^ AML12 cells (each well) were cultured with DMEM/F-12 containing 17.5 mM glucose overnight in XF96 microplates. All reagents were adjusted to pH 7.4. For OCR assays, cells were washed and changed to unbuffered DMEM in the presence of 25 mM glucose and with or without 100 µM ARI and incubated at 37 °C in a non-CO2 incubator for 1 h. For ECAR, cells were washed and replenished with unbuffered DMEM containing non glucose but with or without ARI. Cells were incubated at 37 °C in a non-CO2 incubator for one h before use for the experiments. Three measurements were taken before or after addition of glucose (25 mM), oligomycin (1 µM), 2-Deoxy-D-glucose (2-DG, 150 mM), carbonyl cyanide-4 (trifluoromethoxy) phenylhydrazone (FCCP, 2 µM), and the combination of antimycin A (Ant, 1 µM) and rotenone (Rot, 1 µM). ECAR and OCR data were processed by the Seahorse Wave software.

### Mitochondrial reactive oxygen species (ROS) production as analyzed by MitoSOX Red in AML12 cells

*Ar*-AML12 and NC-AML12 cells were plated and cultured in 6-well plate DMEM/F-12 media. After 25 mM glucose treatment for 4 h, MitoSOX Red (Invitrogen) solution was added at a final concentration 5 µM and the cells were incubated at 37°C for 30 min. Cells were then washed with preheated Hank’s Balanced Salt Solution and digested with trypsin-EDTA for 3-5 min. Cells were collected, washed and passed through a 70 µm filter to prepare single-cell suspensions. The single-cell suspensions were subjected to the flow cytometer (Beckman FC500) and MitoSOX Red fluorescence was detected at 610 nm.

### Metabolomic analyses of AML12 cells

About 8 × 10^5^ *Ar*-AML12 and NC-AML12 cells were plated and cultured in 6-cm dish in normal DMEM/F-12 media until attachment. After 25 mM glucose treatment for 4 h, cells were washed with ice-cold PBS once and then 1 ml pre-cold 80% (v/v) methanol. Cells were collected and homogenized with ultrasonic homogenizer in the ice-cold water bath for 5 min to extract metabolites. The cell lysate was centrifugated at 14,000 ×g for 10 min at 4 °C. The supernatant was dried with the vacuum concentrator (Labconco CentriVap) at 4 °C. The dried supernatant was reconstituted with 100 μL of 50% (v/v) acetonitrile solution and vortexed for 5 minutes before centrifugation at 14,000 ×g for 10 min at 4 °C. The supernatants were subjected to LC-MS/MS (AB SCIEX QTRAP 5500) for analysis.

The liquid chromatography with SCIEX ExionLC AD was prepared and all chromatographic separations were performed with a Millipore ZIC-pHILIC column (5 μm, 2.1× 100 mm internal dimensions, PN: 1.50462.0001). The column was maintained at 40 °C and the injection volume of all samples was 2 μL. The mobile phase consisted of 15 mM ammonium acetate and 3 ml/L ammonium hydroxide (> 28%) in LC-MS grade water (mobile phase A) and LC-MS grade 90% (v/v) acetonitrile-HPLC water (mobile phase B) run at a flow rate of 0.2 mL/min. The analysts were separated with the following gradient program: 95% B held for 2 min, increased to 45% B in 13 min, held for 3 min, and the post time was set for 4 min.

The QTRAP 5500 mass spectrometer (AB SCIEX) using an Turbo V ion source. The ion source was run in a negative mode (spray voltage of -4,500 V) and a positive mode (spray voltage of 5,500 V), with Gas-1 50 psi and Gas-2 55 psi and Curtain gas 35 psi.

Metabolites were measured using the multiple reactions monitoring mode (MRM) optimized with analytical standards. All data were collected using Analyst software (AB SCIEX) and the relative amounts of metabolites were analyzed using MultiQuant software (AB SCIEX). In case the same metabolite was detected by both positive ion mode and negative ion mode, the one with higher signal level was selected, unless otherwise indicated. The processed MS data were eventually analyzed with MetaboAnalyst 5.0 (https://www.metaboanalyst.ca/).

### Other assays

Please see the Supplementary Information for other assays.

### Statistical analyses

All data were expressed either as the mean ± SD or as the mean ± SEM. Students’ *t*-test was used for pair-wise comparisons and One-way ANOVA with Multiple Comparison for multi-group analyses. Probability values less than 0.05 were considered to be statistically significant and those less than 0.01 very significant and those less than 0.001 highly significant.

## Supporting information

Song_et_al_2024_Supplementary Information

## ACKNOWLEDGEMENTS

This work was supported in part by Chinese 973 Program project #2015CB553804, National Natural Science Foundation of China (Grant No. 32160165), Natural Science Foundation of Tibet Autonomous Region (Grant No. XZ202201ZR0065G).

The authors thank other members of Yang lab for their assistance in certain experiments.

## AUTHOR CONTRIBUTION

JYY conceived and initiated the project. FZ, WL, and JYY designed and supervised the experiments. DS performed the animal studies and cell culture analyses. DY performed the Seahorse analyses and the metabolomic studies. FZ, WL, DS, DY and JYY analyzed the data and reviewed the manuscript. JYY wrote the manuscript.

