## Supplementary material for "Hepatic aldose reductase drives a Warburg effect-like metabolic reprogramming to promote insulin resistance, fatty liver and obesity": Song_et_al_2024_Supplementary Information

1. ***Supplemental Materials and Methods***

***Reagents.*** PCR primers (Table S1), antibodies (Table S2) and cell media and other reagents (Table S3) used in this study were listed with catalog numbers and the manufacturers/suppliers. Ar inhibitor zopolrestate (ARI) was a kind gift from the Pfizer Inc. PCR primers were provided by Sangon, Shanghai, China.

***Plasmids and lentiviruses.*** Mouse Ar overexpressing pFlag-mAr and lentivirus-mediated *Ar* knockdown plasmid LV-shAr-1 and LV-shAr-2 and the controls were constructed previously by our lab [1,2].

Plasmids for lentiviral vector construction and virus package VSVG, pMDL, REV and pLV-TagII were kind gifts from Prof. Jiahuai Han, Xiamen University. To prepare stable cells overexpressing mouse Ar, Mouse Ar cDNA fragment was amplificated from pFlag-mAR plasmid and sub-cloned into pLV-TagII plasmid by ligation-independent cloning to generate pLV-TagII-mAR. The *Ar*-overexpressing lentivirus and the control lentivirus were generated by co-transfection with plasmids VSVG, pMDL, REV and pLV-TagII or pLV-TagII-mAR in HEK293T cells, respectively. After 24 hours, the supernatants containing the viral particles were collected and passed through a 0.45 um filter. The viral particles can be either stored at -80 ºC or use immediately.

***Quantitative analyses of mRNA expression by real-time PCR and western blot analysis of proteins.*** Total RNA was isolated from AML12 cells or tissue using Trizol Reagent (Invitrogen) according to the manufacturer's protocol. DNA amplification was carried out using a High Fidelity primeScript TM RT-PCR Kit (Takara) and the expression of mRNAs were analyzed by Real-time PCR using primers listed on Table S1. Western blots were performed according to standard protocols using antibodies listed on Table S2. The detection was achieved using the Immobilon Western Chemiluminescent HRP Substrate kit (Millipore, Massachusetts, USA).

***Nile red staining of glucose fed AML12 cells.*** *Ar*-overexpressing AML12 cell (*Ar*-AML12) and its control (NC-AML12) were cultured in DMEM containing 5.6 mM glucose for 14 h. Cells were then cultured in media containing either 25 mM glucose or 25 mM glucose + 100 μM ARI for another 24 h, respectively. Then cells were fixed in 3% paraformaldehyde fix solution, stained with Nile red (0.05 μg/ml) (Sigma, Cat # N3013) for 10 min and counter-stained with 4’,6-diamidino-2-phenylindole (DAPI), then washed three times with PBS before being visualized by confocal laser scanning microscopy. Images were obtained using a ZEISS LSM 900 confocal microscope with ZEN software (Carl Zeiss GmbH, Jena, Germany).

***$NADP^+$/NADPH measurements.*** About 1 × $10^6$ *Ar*-AML12 and NC-AML12 cells were seeded onto the 6-well plates and grown in DMEM/F-12 media. On the day of the experiments, cells were treated with 25 mM glucose for 4 h, with or with no ARI. Then cells were washed with 1 × PBS and lysed with $NADP^+$/NADPH extraction buffer directly. $NADP^+$/NADPH was assayed according to the manufacturer's protocol ($NADP^+$/NADPH Assay, Cat # S0179, Beyotime, Shanghai, China).

***2-deoxy-glucose (2-DG) uptake***. About 1 × $10^6$ Ar-AML12 and NC-AML12 cells were seeded into the 6-well plate and cultured overnight. The next day, the medium was changed with fresh one containing neither FBS and nor glucose for 1 h. Then the medium was discarded and treated with new medium containing FBS and 2-DG (25 mM) for 10 min. The cells were washed with ice-cold mannitol (5% in water) immediately and then added with methanol (80% in water) at -80 ºC. 2-DG was detected with LC-MS as described in the main text, reading only 2-DG and the internal standards.

***DNL assays in Ar deficient or Ar inhibitor treated mice***. For blood TG assays, male and female $Ar^{-/-}$ C57Bl/6 mice and wildtype controls were bred and weaned normally. On the day they reached the age of eight weeks, food was removed starting at 10:00 am in the morning. Four hours later, blood TG levels were measured and recorded as the initial levels. The mice were then intraperitoneally injected with a glucose solution at the dosage of 4 g/kg body weight. After one hour, blood TGs were measured and recorded.

For isotopic TG tracing, male wildtype C57BL/6 (WT) were bred and weaned normally. Three days before they reached the age of 8 wk, they were randomly divided into two groups. One group was pretreated with a chow supplemented with an aldose reductase inhibitor (ARI) zopolrestat for three days while the other received regular chow. The ARI chow was prepared by mixing the regular chow with zopolrestat powder to obtain a targeted dosage of 50 mg/kg [3,4] based on a daily chow consumption of 4 g /mouse, which was determined from the results of preliminary feeding experiments. Following the three-day pretreatment, both groups of mice were fasted for 4 h starting at 10:00 am in the morning on the 4th day. At 14:00 pm, the mice were intraperitoneally injected with a glucose solution that contains $^{14}C$-labeled glucose. The glucose solution was so prepared that each mouse received around 40 μCi of U-$^{14}C$-glucose (CFB96, 291 mCi/mmol, Amersham, New Jersey) while glucose loading was at the dosage of 2 g/kg body weight. Exactly one hour later, the mice were sacrificed and the liver tissues were rapidly dissected and frozen immediately in liquid nitrogen. Total lipids were extracted with chloroform/methanol (2:1) from liver according to Folch et al. [5]. Lipid samples were separated on silica gel thin layer chromatography plates (TLC Silica Gel G60, Merck, Darmstadt, Germany) using petroleum ether-diethyl ether-acetic acid (80:30:1) as the developing solvent. A non-radioactive lipid sample extracted from the epididymal fat tissues of normal mice was prepared and used to load and run along with the samples of interest. This standard sample therefore not only served as a reference for TGs but also was used to construct a standard curve for TG quantitation in every TLC run. The lipids were visualized by spraying the TLC plates with 10% cupric sulfate in 8% phosphoric acid and baking in a vacuum oven. The TLC plates were scanned and the lipid spots were digitally quantitated with SigmaScan Pro Version 5.0.0 (SPSS Inc., Chicago, Illinois). TG spots were carefully scraped off and the radioactivity was measured by liquid scintillation counting on a Beckman LS 6500 scintillation counter (Fullerton, CA). The counts per minute were corrected with counting efficiency to disintegrations per minute. The yields of both total lipids and $^{14}C$-labeld TGs were comparable to that from rat liver [6].

***Liver tissue histology***. For histological analyses, 130 days Agouti mice and the controls or 154 days *Ar* overexpressing transgenic mice and the controls were sacrificed and liver tissues were rapidly dissected and fixed in 10% buffered formalin (v/v) for 24 h before being embedded in paraffin, respectively. Liver tissue sections at 5-μm thickness.

For Liver H & E staining, sections were first dipped in haematoxylin (Jiancheng Biotech, Nanjing, China) and wash with water, then incubated with eosin and washed thoroughly and covered with mounting medium, the sections were examined under a light microscope.

For Oil red O staining, sections of liver were quickly frozen in liquid nitrogen, embedded in optimal cutting temperature compound (Sakura Finetek, Inc., Torrance, CA, USA), cut into 10-μm sections, mounted on glass slides and Oil red O stained (Jiancheng Biotech, Nanjing, China) according to the manufacturer's protocol.

***Lactate secretion in AML12 cells.*** About 1 × $10^6$ *Ar*-AML12 and NC-AML12 cells were seeded onto the 6-well plates and grown in DMEM/F-12 containing 17.5 mM glucose. On the day of the experiments, cells were washed with 1× PBS twice and then fresh media with or without 100 μM ARI were added. The medium supernatants were collected at 2, 4, and 8 h. Lactate contents were assayed according to the manufacturer's protocol (Lactic Acid assay Kit, Cat # A019-2-1, Jiancheng, Nanjing, China).

***Blood or serum sample analyses.*** Blood samples were collected by eye ball puncture and placed at room temperature for 2 hours then centrifuged at 4 °C, 2000 rpm × 20 min to get the serum. Blood or serum biochemical parameters including serum insulin, leptin, fructose TG cholesterol, alanine aminotransferase and liver fructose, liver TG and liver cholesterol were assayed by reagent kits or devices as listed on Table S2, according to the instructions of the manufacturers.

***Glucose and insulin tolerance tests and HOMR-IR.*** For the glucose tolerance tests (GTT) were performed by intraperitoneal injection of mice with glucose (2.0 g/kg body weight) after overnight fasting for 16 h. Glucose in a tail vein blood sample was measured at 0-, 15-, 30-, 60-, 120-min using a Glucometer (OneTouch Ultra, LifeScan, Milpitas, CA, USA). The insulin tolerance tests (ITT) was performed by intraperitoneal injection of insulin (0.75 U/kg body weight) 6 h after food withdrawal [7]. Glucose in a tail vein blood sample was measured at 0-, 15-, 30-, 60-, 120-min using the Glucometer. HOMA-IR was calculated by the formula: fasting plasma glucose (mmol/L) × fasting insulin (mIU/L) / 22.5.

***Supplemental Figure and Table Legends***

***Fig. S1 Related to Introduction*. *The conventional glycolysis and the pseudo-glycolysis*.** DHAP, dihydroxyacetone phosphate; F1P, fructose 1-phosphate; F6P, fructose 6-phosphate; FBP, fructose 1,6 bisphosphate; G6P, glucose 6-phosphate; GA, glyceraldehyde; GA3P, glyceraldehyde 3-phosphate.

***Fig. S2 Related to Fig. 2. Metabolomic analyses of the control and stable Ar-overexpressing AML12 cells.*** The LC-MS metabolomic data were analyzed by MetaboAnalyst 5.0 ($n$ = 3). NC-AML12 cells, normal AML12 cells; Ar-AML12, stable *Ar*-overexpressing AML12 cells. ***A***. PCA scores 2D-plot of based on 280 detected metabolites. ***B***. NADP+/NADPH ratio as determine by spectrophotometer at OD450 ($n$ = 5-6)**.** Values were expressed as the mean ± SD. NS, not significant; *, $p < 0.05$; **, $p < 0.01$; ***, $p < 0.001$. ***C***. Variable importance in projection scores as analyzed by the partial least squares discriminant analysis (PLS-DA, 280 detected metabolites).

***Fig. S3 Related to Fig. 2. Metabolomic analyses of the control and stable Ar-overexpressing AML12 cells*** ($n$ = 3)**.** ***A.*** Heatmap of enriched pyrimidine and purine derivatives. ***B.*** Heatmap of metabolites enriched in the galactose metabolism pathway. ***C.*** Heatmap of metabolites enriched in the DNL pathway. GO3P, glycerol 3-phosphate. ***D***. Heatmap of reduced glutathione (GSH) and oxidized glutathione (glutathione disulfide, GSSG) as detected by the positive or negative ions respectively.

***Fig. S4 Related to Fig. 2. Metabolomic analyses of the control and stable Ar-overexpressing AML12 cells*** ($n$ = 3)**.** The abundancy of 27 selected metabolites as assayed by LC-MS. The scales were in arbitrary unit.

***Fig. S5 Related to Fig. 3. Irs-1 and Lkb1-Ampk signaling in Ar overexpressing or knockdown AML12 cells.*** Values were expressed as the mean ± SEM. NS, not significant; ***, $P < 0.001$. Cells were transfected with pFlag-CMV2/pFlag-mAr for 24 h or infected with lentiviruses LV-CK or LV-shAr-1/LV-shAr-2 for 96 h. Cells were collected for mRNA analyses by qPCR using 18S ribosomal RNA as a calibrator. Experiments were performed with at least three separate samples analyzed in triplicate (*n* = 3). ***A.*** *Ar* mRNA expression in *Ar* overexpressing or *Ar* knockdown AML12 cells. ***B***. *Irs1* mRNA expression in *Ar* overexpressing or *Ar* knockdown AML12 cells.

***Fig. S6 Related to Fig. 5. Construction of liver-specifically human AR-overexpressing transgenic mice and hepatic protein expression of Irs1, Lkb1, Ampkα and Acc. A.*** The design of the liver-specific *AR* overexpressing transgene. A 951-bp human *AR* cDNA derived from human LO2 cells was fused with the Flag protein tag fragment. This Flag-tagged *AR* was then inserted into the plasmid LIV-Le6 at ClaI/XhoI sites such that the expression of human *AR* cDNA is under the transcriptional control of the *apoE* promoter. ***B***. *AR* mRNA and protein expression in the livers and kidney tissues of transgenic and control mice after weaning. ***C***. Hepatic transgene expression in the transgenic and control mice after weaning as determined by immunohistochemistry of Flag (brown staining). ***D***. Experimental design for liver-specific *Ar*-transgenics. ***E***. Irs1 protein expression of livers from the transgenic and control mice with or without 10% glucose supplement at the age of 154-d (*n* =3). ***F***. Phosphorylated and unphosphorylated Lkb1, Ampkα and Acc protein expression of livers from the transgenic and control mice with or without 10% glucose supplement at the age of 154-d (*n* =3).

***Fig. S7 Related to Fig. 6. Effects of Ar deficiency biochemical parameters, liver steatosis and chow consumption in Agouti yellow obese mice and the controls.*** Only male mice at the age of 130-d were used. Values were expressed as the mean ± SEM. Pair-wise comparisons were made for $a/a::Ar^{+/+}$ versus $a/a::Ar^{-/-}$ (in grey *) *or* $a/a::Ar^{+/+}$ versus $A^{y}/a::Ar^{+/+}$ (in yellow #) or $A^{y}/a::Ar^{+/+}$ versus $A^{y}/a::Ar^{+/+}$ (in yellow *). NS, not significant; * and #, $p < 0.05$; ** and ##, $p < 0.01$; ###, $p < 0.001$; ####, $p < 0.0001$. ***A***. Experimental design for 4 groups of mice. ***B-G***, Biochemical parameters in 4 groups of mice (*n* = 3-8), blood glucose (***B***), liver fructose (***C***) and serum levels of insulin (***D***), leptin (***E***), fructose (***F***), TG (***G***). ***H.*** Hepatic steatosis as analyzed by Oil-red O staining of livers of 4 groups of mice. Magnification, 200×. Data were representative of at least three mice (*n* = 3). ***I.*** Daily chow intake of $A^{y}/a::Ar^{+/+}$ and $A^{y}/a::Ar^{-/-}$ at the age of 70-d. Values were expressed as the mean ± SEM. NS, not significant, *n*

= 4-5.

***Table S1 PCR primers used.***

***Table S2 PCR Antibodies used.***

***Table S3 Cell media and other reagents used.***

2. ***Supplemental Figures and Tables***

***Fig. S1***

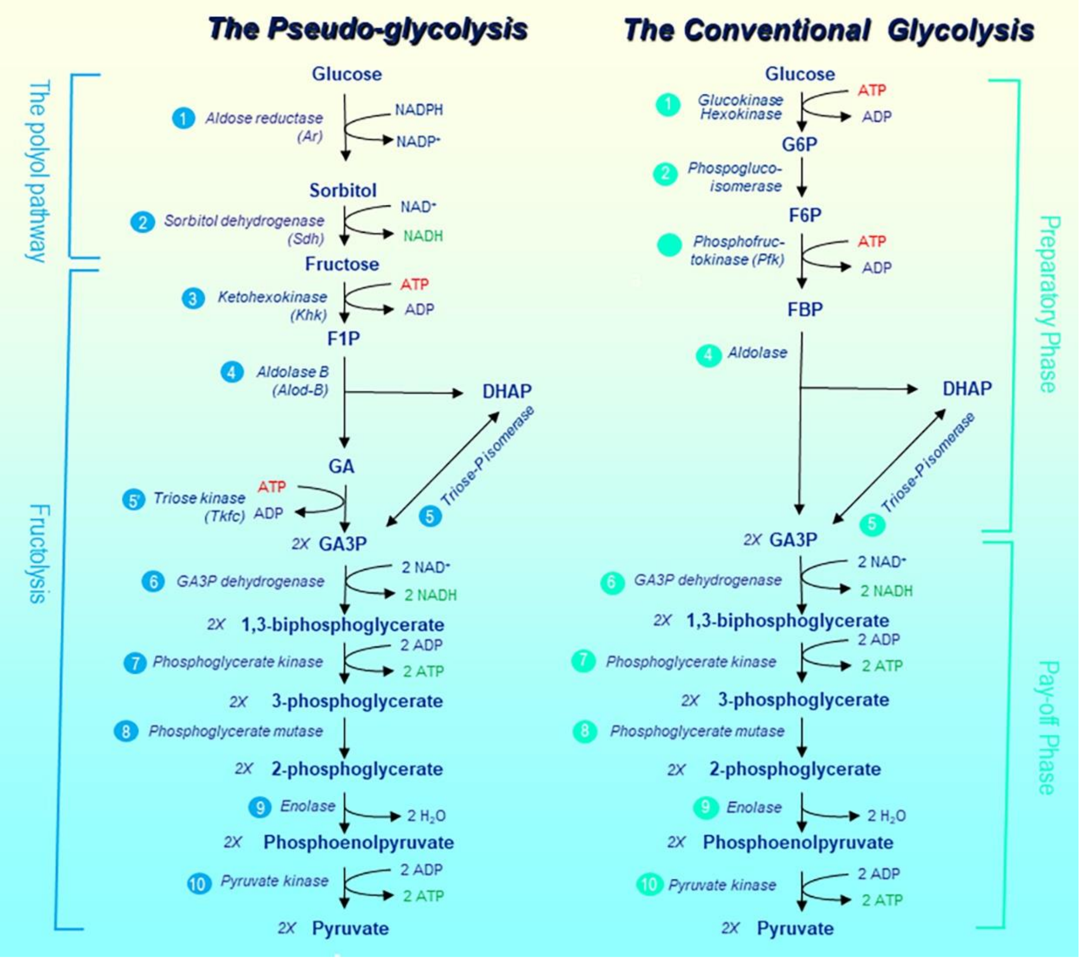

*Fig. S2*

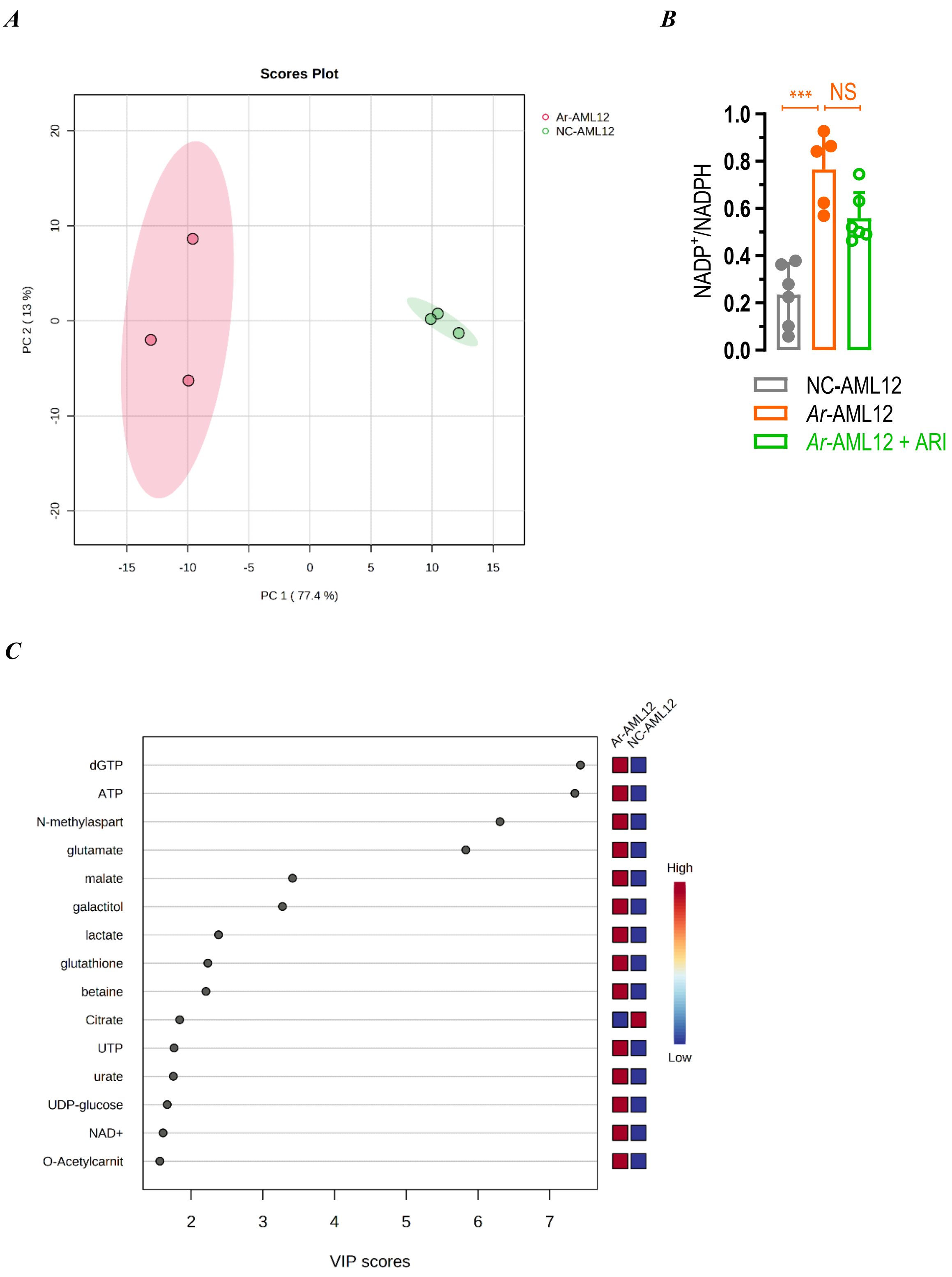

A
B
Scores Plot
Ar-AML12
NC-AML12
PC 2 ( 13 %)
PC 1 ( 77.4 %)
-15
-10
-5
0
5
10
15
20
10
0
-10
-20
***
NS
1.0
0.8
0.6
0.4
0.2
0.0
NADP+/NADPH
NC-AML12
Ar-AML12
Ar-AML12 + ARI
C
Ar-AML12
NC-AML12
dGTP
ATP
N-methylaspart
glutamate
malate
galactitol
lactate
glutathione
betaine
Citrate
UTP
urate
UDP-glucose
NAD+
O-Acetylcarnit
High
Low
2
3
4
5
6
7
VIP scores

*Fig. S3*

A

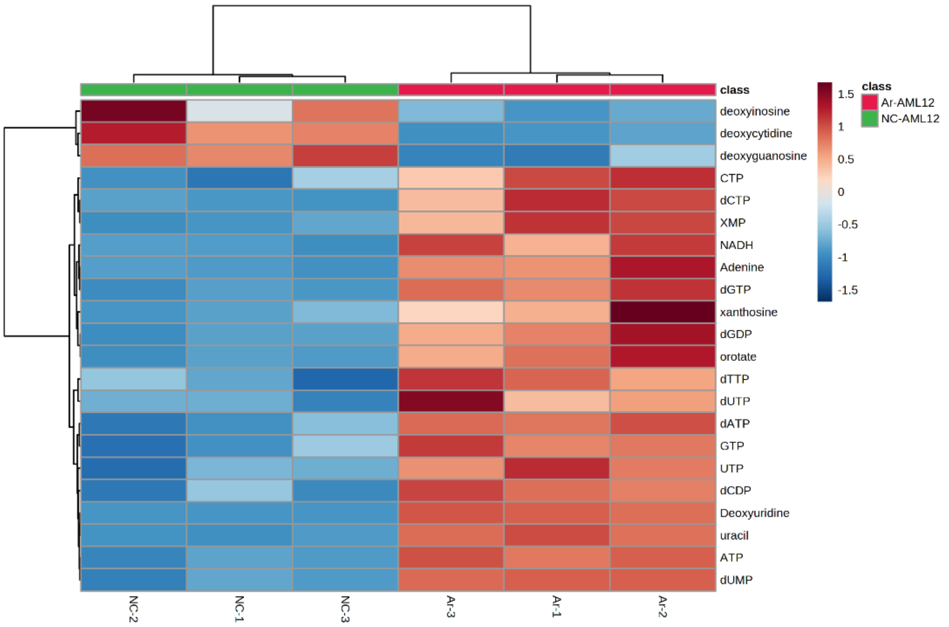
class
deoxyinosine
deoxycytidine
deoxyguanosine
CTP
dCTP
XMP
NADH
Adenine
dGTP
xanthosine
dGDP
orotate
dTTP
dUTP
dATP
GTP
UTP
dCDP
Deoxyuridine
uracil
ATP
dUMP
NC-2
NC-1
NC-3
Ar-3
Ar-1
Ar-2
1.5
1
0.5
0
-0.5
-1
-1.5
class
Ar-AML12
NC-AML12

B

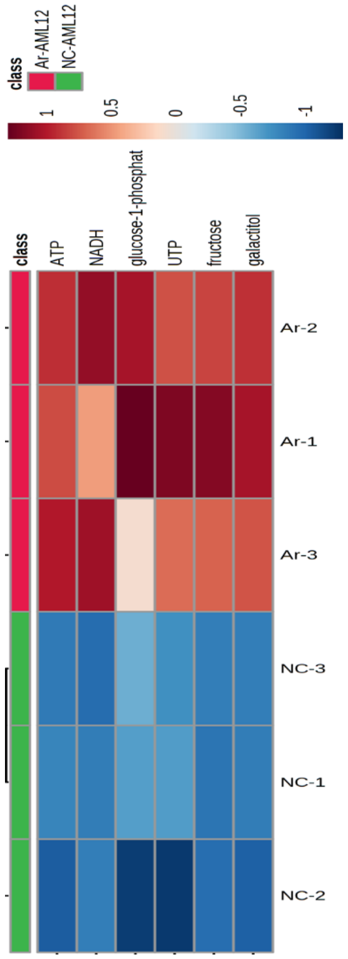
class
Ar-AML12
NC-AML12
1
0.5
0
-0.5
-1
class
ATP
NADH
glucose-1-phosphat
UTP
fructose
galactitol
Ar-2
Ar-1
Ar-3
NC-3
NC-1
NC-2

C

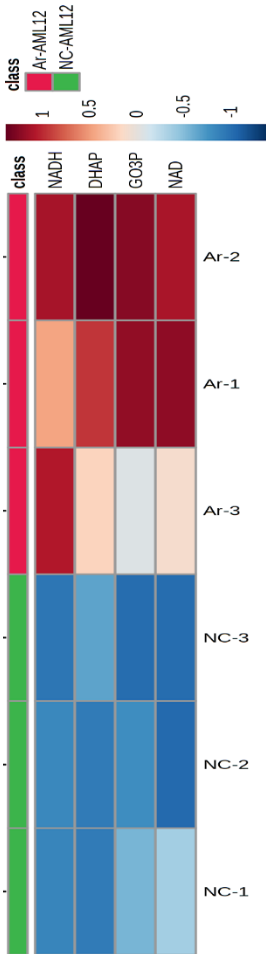
class
Ar-AML12
NC-AML12
1
0.5
0
-0.5
-1
class
NADH
DHAP
GO3P
NAD
Ar-2
Ar-1
Ar-3
NC-3
NC-2
NC-1

D

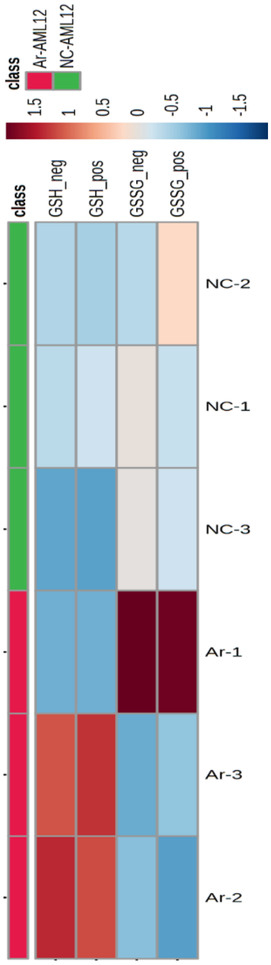
class
Ar-AML12
NC-AML12
1.5
1
0.5
0
-0.5
-1
-1.5
class
GSH_neg
GSH_pos
GSSG_neg
GSSG_pos
NC-2
NC-1
NC-3
Ar-1
Ar-3
Ar-2

*Fig. S4*

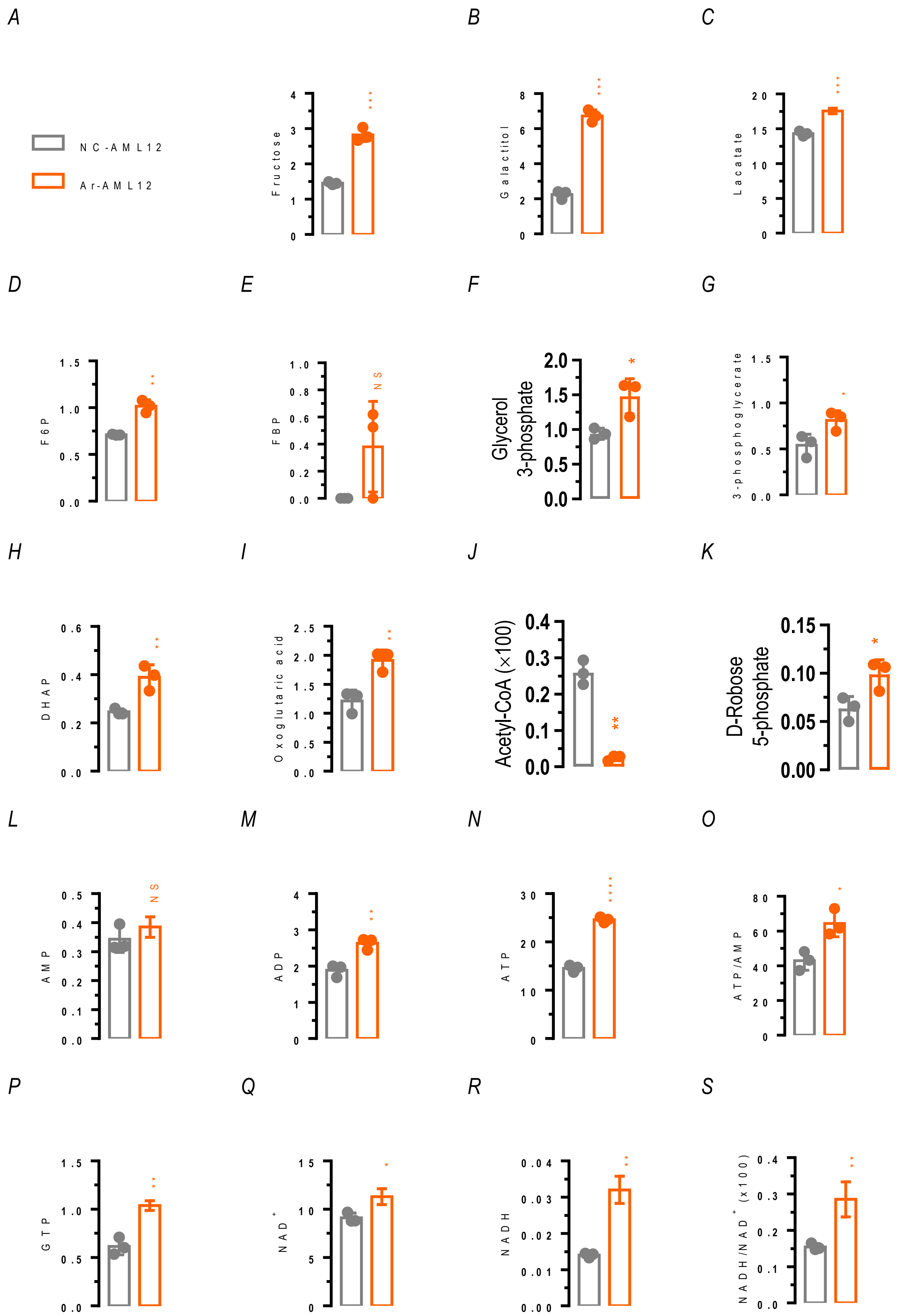
A
B
C
NC-AML12
Ar-AML12
Fructose
Galactitol
Lacatate
D
E
F
G
F6P
FBP
NS
Glycerol 3-phosphate
3-phosphoglycerate
H
I
J
K
DHAP
Oxoglutaric acid
Acetyl-CoA (×100)
D-Robose 5-phosphate
L
M
N
O
AMP
NS
ADP
ATP
ATP/AMP
P
Q
R
S
GTP
NAD+
NADH
NADH/NAD+ (x100)

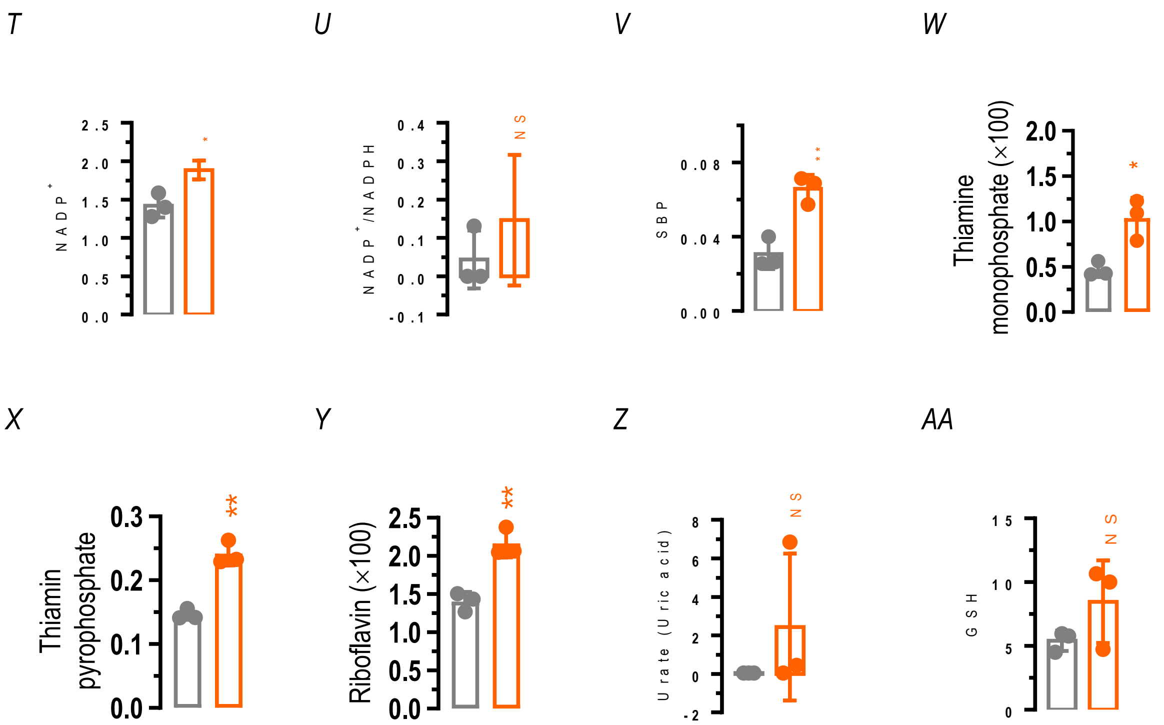

T
U
V
W
NADP+
NADP+/NADPH
SBP
Thiamine monophosphate (×100)
*
NS
**
*
X
Y
Z
AA
Thiamin pyrophosphate
Riboflavin (×100)
Urate (Uric acid)
GSH
**
**
NS
NS

***Fig. S5***

A

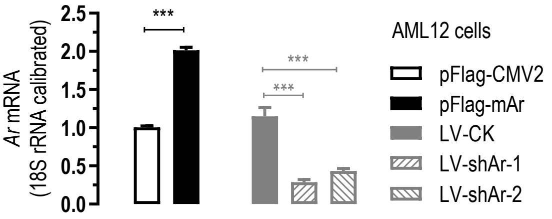

Ar mRNA
(18S rRNA calibrated)
2.5
2.0
1.5
1.0
0.5
0.0
***
***
***
AML12 cells
pFlag-CMV2
pFlag-mAr
LV-CK
LV-shAr-1
LV-shAr-2


B

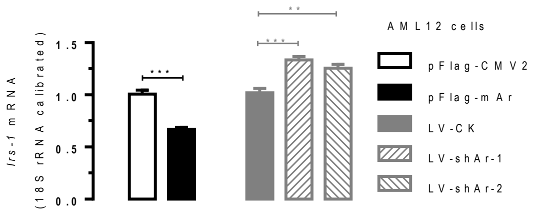

Irs-1 mRNA
(18S rRNA calibrated)
1.5
1.0
0.5
0.0
***
**
***
AML12 cells
pFlag-CMV2
pFlag-mAr
LV-CK
LV-shAr-1
LV-shAr-2

*Fig. S6*

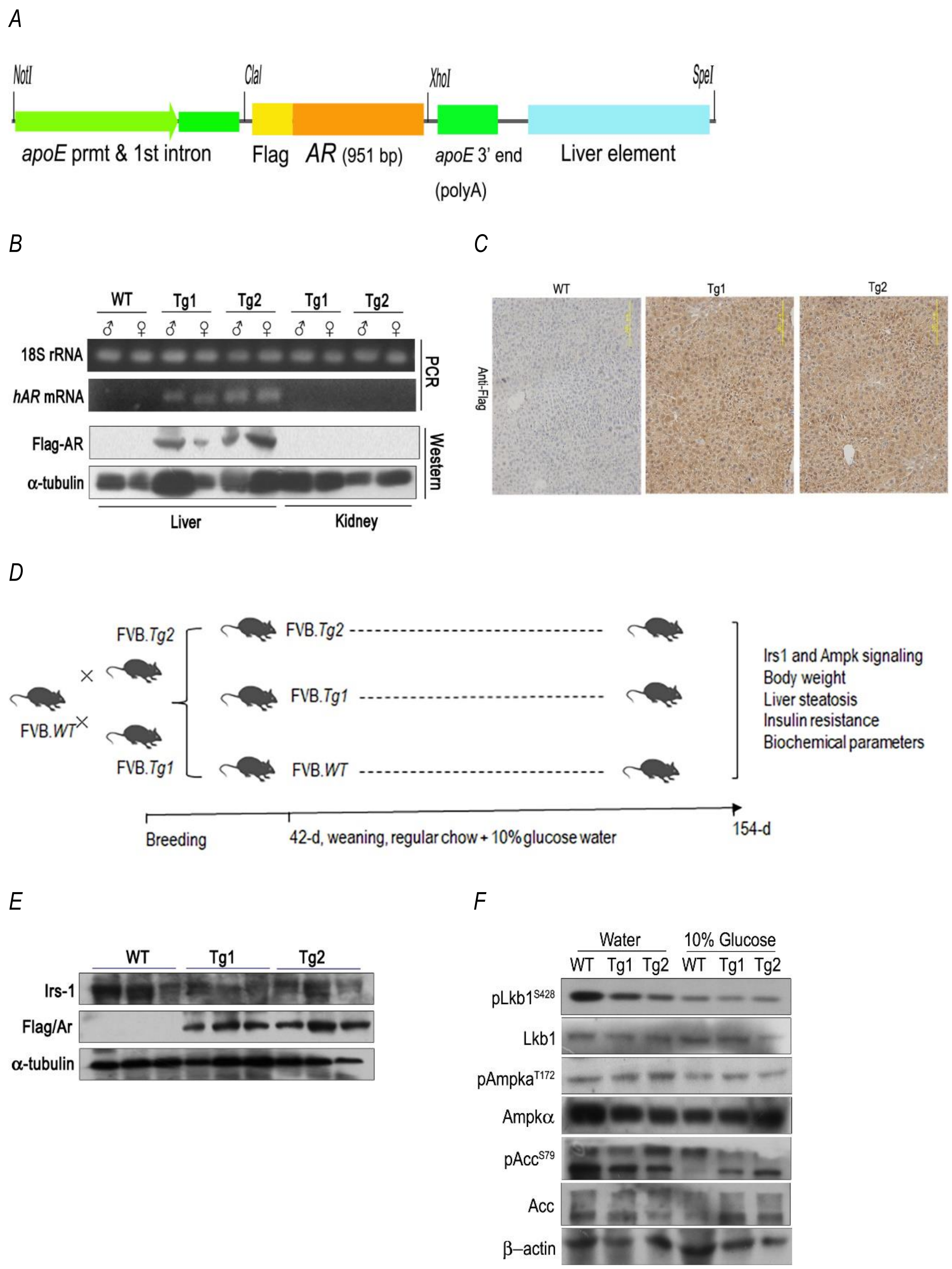
A
NotI
ClaI
XhoI
SpeI
apoE prmt & 1st intron
Flag
AR (951 bp)
apoE 3' end
(polyA)
Liver element
B
WT Tg1 Tg2 Tg1 Tg2
♂ ♀ ♂ ♀ ♂ ♀ ♂ ♀ ♂ ♀
18S rRNA
hAR mRNA
PCR
Flag-AR
α-tubulin
Western
Liver
Kidney
C
WT
Tg1
Tg2
Anti-Flag
D
FVB.Tg2
FVB.Tg1
FVB.WT
Irs1 and Ampk signaling
Body weight
Liver steatosis
Insulin resistance
Biochemical parameters
Breeding
42-d, weaning, regular chow + 10% glucose water
154-d
E
WT Tg1 Tg2
Irs-1
Flag/Ar
α-tubulin
F
Water
10% Glucose
WT Tg1 Tg2 WT Tg1 Tg2
pLkb1S428
Lkb1
pAmpkaT172
Ampkα
pAccS79
Acc
β–actin

*Fig. S7*

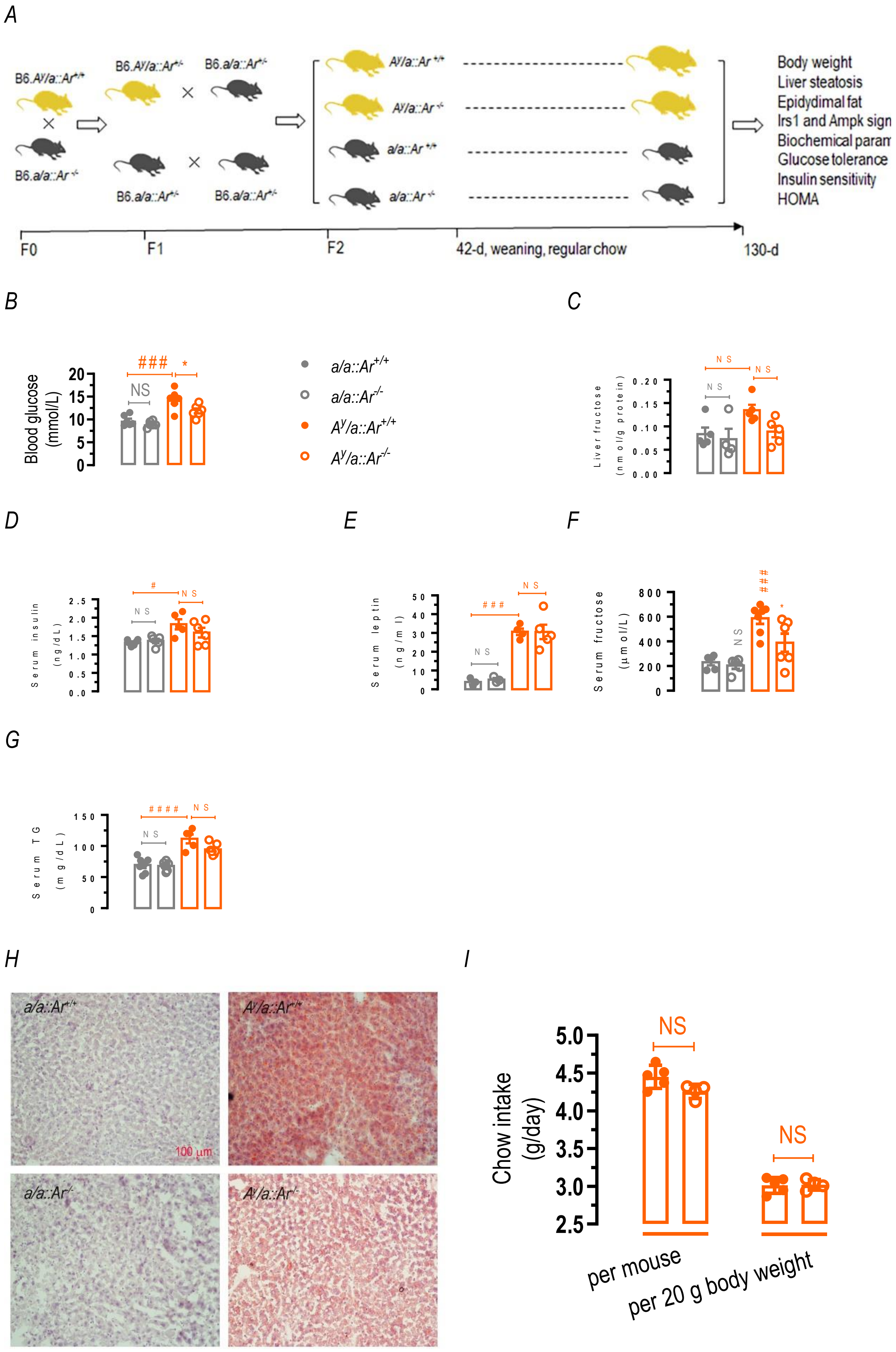

A
B6.A^y/a::Ar^+/+
B6.a/a::Ar^+/-
B6.A^y/a::Ar^+/-
B6.a/a::Ar^+/-
A^y/a::Ar^+/+
A^y/a::Ar^-/-
a/a::Ar^+/+
a/a::Ar^-/-
Body weight
Liver steatosis
Epidydimal fat
Irs1 and Ampk sign
Biochemical param
Glucose tolerance
Insulin sensitivity
HOMA
F0
F1
F2
42-d, weaning, regular chow
130-d
B
Blood glucose (mmol/L)
C
Liver fructose (nmol/g protein)
D
Serum insulin (ng/dL)
E
Serum leptin (ng/ml)
F
Serum fructose (μmol/L)
G
Serum TG (mg/dL)
H
100 μm
I
Chow intake (g/day)
per mouse
per 20 g body weight
NS

***Table S1***

| Gene | Gene ID | Primer sequence (from 5' to 3') | Product length |
|---|---|---|---|
| Flag-hAR | Transgene | Forward: ACTGTGCCCATGTGTACCAGA<br>Reverse: CCAATAGCTTTCACCAGCCCT | 343 bp |
| *Akr1b3* *(Ar)* | NM_009658.3 | Forward: AACAGGAACTGGAGGGTGTGC<br>Reverse: ACTTGAGGAGGAGCAGGGATC | 110 bp |
| *Irs-1* | NM_010570.4 | Forward: TGAGCCCCAAGAGTGTATCTG<br>Reverse: CAATGTCAGGGGAGCAACTAC | 123 bp |
| 18S rRNA | NR_003286 | Forward: CGACGACCCATTCGAACGTCT<br>Reverse: CTCTCCGGAATCGAACCCTGA | 103 bp |

*Table S2*

| Antibody | Catalog number | Company |
|---|---|---|
| Irs1 | 06-248 | Upstate, Massachusetts, USA |
| Ar | AB60849a | Sangon Biotech, China |
| Flag M2 | F1804 | Sigma, St. Louis, USA |
| phospho-Ser-428 Lkb1 | ab63473 | Abcam, Cambridge, UK |
| Lkb1 | 3047 | Cell Signaling, Danvers, USA |
| phospho-Thr-172 Ampkα | 2535 | Cell Signaling |
| Ampkα | 2532 | Cell Signaling |
| phospho-Ser-79 Acc | 07-303 | Millipore, Massachusetts, USA |
| Acc | 3676 | Cell Signaling |
| β-actin | A1978 | Sigma, Shanghai, China |
| α-tubulin | sc5286 | Santa Cruz, Dallas, USA |
| β-tubulin | MA8281 | Abmart, Shanghai, China |
| HRP-second antibody | 31430 | Pierce, Rockford, USA |

*Table S3*

| Item name | Cat # | Supplier |
| --- | --- | --- |
| 2-deoxy glucose (2-DG) | A602241 | Sangon Biotech, Shanghai, China |
| Antimycin A | A8674 | Sigma, Shanghai, China |
| Blood TG | Accutrend® GCT system | Boehringer Mannheim, Germany |
| DMEM/F-12 medium | 12400-024 | GIBCO, Grand Island, NY, USA |
| DMEM (high glucose) | 12800-017 | GIBCO |
| DMEM (low glucose) | 31600-034 | GIBCO |
| Glucose test strips | 01-0228 | Onetouch Ultra, Lifescan, CA |
| Glucometer Elite | Model 3049C | Bayer, USA |
| EnzyChrom™ Fructose Assay Kit | EFRU-100 | Bio Assay System, California, USA |
| FCCP | S8276 | Selleck, Shanghai, China |
| Fetal bovine serum (FBS) | | Sijiqing, Hangzhou, China |
| H&E | D006-1-1 | Jiancheng Biotech, Nanjing |
| Lactic acid Assay Kit | A019-2-1 | Jiancheng Biotech, Nanjing |
| MitoSOX Red | M36008 | Invitrogen, Shanghai, China |
| Mouse Insulin ELISA kit | E0310004 | Blue Gene Biotech Co., Ltd, Shanghai, China |
| Mouse Leptin ELISA Kit | 90030 | CRYSTAL CHEM INC., Downers Grove, USA |
| $NADP^+$/NADPH | S0179 | Beyotime Biotech, Shanghai, China |
| Nile red | N3013 | Sigma |
| OCT | | Sakura Finetek, Inc, Torrance, CA, USA |
| Oligomycin | S1478 | Selleck, Shanghai, China |

| | | |
|---|---|---|
| Oil red O | O0625 | Sigma, Shanghai, China |
| Polybrene | H9268 | Sigma |
| Rotenone | S2348 | Selleck |
| Serum TG reagent kit | F001-2 | Jiancheng Biotech, Nanjing, China |
| Serum alanine aminotransferase (ALT) reagent kit | 001020 | BHKT, Beijing, China |
| Total serum cholesterol reagent kit | F002-2 | Jiancheng Biotech, Nanjing |

### 4. *Supplemental References*


1. Qiu, L., Wu, X., Chau, J.F., Szeto, I.Y.Y., Tam, W.Y., Guo, Z., Chung, S.K., Oates, P.J., Chung, S.S.M., and Yang, J.Y. (2008). Aldose reductase regulates hepatic peroxisome proliferator-activated receptor alpha phosphorylation and activity to impact lipid homeostasis. J. Biol. Chem *283*, 17175-17183.
2. Qiu, L., Lin, J., Xu, F., Gao, Y., Zhang, C., Liu, Y., Luo, Y., and Yang, J.Y. (2012). Inhibition of aldose reductase activates hepatic peroxisome proliferator-activated receptor-alpha and ameliorates hepatosteatosis in diabetic db/db mice. Exp Diabetes Res *2012*, 789730. 10.1155/2012/789730 [doi].
3. Beyer-Mears, A., Mistry, K., Diecke, F.P., and Cruz, E. (1996). Zopolrestat prevention of proteinuria, albuminuria and cataractogenesis in diabetes mellitus. Pharmacology *52*, 292-302.
4. Tracey, W.R., Magee, W.P., Ellery, C.A., MacAndrew, J.T., Smith, A.H., Knight, D.R., and Oates, P.J. (2000). Aldose reductase inhibition alone or combined with an adenosine A(3) agonist reduces ischemic myocardial injury. Am. J. Physiol Heart Circ. Physiol *279*, H1447-H1452.
5. Folch, J., Lees, M., and Sloane-Stanley, G.H. (1951). A simple method for the isolation and purification of total lipides from animal tissues. J. Biol. Chem *226*, 497-509.
6. Maruhama, Y., and Macdonald, I. (1973). Incorporation of orally administered glucose-U-14C and fructose-U-14C into the triglyceride of liver, plasma, and adipose tissue of rats. Metabolism *22*, 1205-1215.
7. Wei, J., Lin, Y., Li, Y., Ying, C., Chen, J., Song, L., Zhou, Z., Lv, Z., Xia, W., Chen, X., and Xu, S. (2011). Perinatal exposure to bisphenol A at reference dose predisposes offspring to metabolic syndrome in adult rats on a high-fat diet. Endocrinology *152*, 3049-3061. 10.1210/en.2011-0045.
